# Metagenomic Covariation Along Densely Sampled Environmental Gradients in the Red Sea

**DOI:** 10.1101/055012

**Authors:** Luke R. Thompson, Gareth J. Williams, Mohamed F. Haroon, Ahmed Shibl, Peter Larsen, Joshua Shorenstein, Rob Knight, Ulrich Stingl

## Abstract

Oceanic microbial diversity covaries with physicochemical parameters. Temperature, for example, explains approximately half of global variation in surface taxonomic abundance. It is unknown, however, whether covariation patterns hold over narrower parameter gradients and spatial scales, and extending to mesopelagic depths. We collected and sequenced 45 epipelagic and mesopelagic microbial metagenomes on a meridional transect through the eastern Red Sea. We asked which environmental parameters explain the most variation in relative abundances of taxonomic groups, gene ortholog groups, and pathways—at a spatial scale of <2000 km, along narrow but well-defined latitudinal and depth-dependent gradients. We also asked how microbes are adapted to gradients and extremes in irradiance, temperature, salinity, and nutrients, examining the responses of individual gene ortholog groups to these parameters. Functional and taxonomic metrics were equally well explained (75-79%) by environmental parameters. However, only functional and not taxonomic covariation patterns were conserved when comparing with an intruding water mass with different physicochemical properties. Temperature explained the most variation in each metric, followed by nitrate, chlorophyll, phosphate, and salinity. That nitrate explained more variation than phosphate suggested nitrogen limitation, consistent with low surface N:P ratios. Covariation of gene ortholog groups with environmental parameters revealed patterns of functional adaptation to the challenging Red Sea environment: high irradiance, temperature, salinity, and low nutrients. Nutrient acquisition gene ortholog groups were anticorrelated with concentrations of their respective nutrient species, recapturing trends previously observed across much larger distances and environmental gradients. This dataset of metagenomic covariation along densely sampled environmental gradients includes online data exploration supplements, serving as a community resource for marine microbial ecology.

## Introduction

Microbial communities play a central role in energy flow and carbon and nutrient cycling in the oceans. Shotgun sequencing and analysis of microbial community DNA (metagenomics) is a proven tool for understanding the microbial genomic diversity underlying these processes (DeLong et al., 2006; Dinsdale et al., 2008). Distribution of microbial diversity and biogeochemistry is structured in large part by environmental gradients in light, temperature, oxygen, salinity, and nutrients. Oceanographic surveys spanning such environmental gradients, combining metagenomic sequencing and measurement of continuous environmental variables, are enabling quantitative understanding of microbial communities (Gianoulis et al., 2009; Raes et al., 2011). Global oceanographic surveys have sequenced hundreds of surface and moderately deep (epipelagic and mesopelagic) ocean microbial communities (Rusch et al., 2007; Sunagawa et al., 2015), cataloging the vast genomic diversity of ocean microbes; further analyses of these data have identified correlations between environmental parameters and genetic community traits (gene ortholog groups and pathways) with predictive power (Gianoulis et al., 2009; Raes et al., 2011; Barberán et al., 2012). Local studies at individual ocean sites, meanwhile, have shown how microbial taxa and gene ortholog groups are partitioned at greater detail along the water column and between discrete ocean environments (DeLong et al., 2006; Coleman and Chisholm, 2010; Ghai et al., 2010; Thompson et al., 2013). Depth is a critical factor behind community structure in the open ocean (DeLong et al., 2006), and dense sampling is capable of capturing subtle changes in environmental parameters with sufficient replication for statistical power.

The Red Sea is an ideal oceanic site for dense sampling of metagenomes to study environment-microbe covariation. The Red Sea is a deep (>2000 m) incipient ocean with strong latitudinal and depth-dependent gradients in temperature, salinity, oxygen, and nutrients (Edwards, 1987). Like the open-ocean gyres of the Atlantic and Pacific Oceans, the Red Sea is oligotrophic with surface waters dominated by the picoplankton *Prochlorococcus* and *Pelagibacter* (Ngugi and Stingl, 2012). More so than these open-ocean gyres, however, the Red Sea lies at pelagic extremes of irradiance, temperature, and salinity. The Red Sea experiences a late-summer southern influx of water called the Gulf of Aden Intermediate Water (GAIW), a foreign water mass that is cooler, fresher, and more nutrient-rich than the native Red Sea water mass. The Red Sea is compact enough to sample across these gradients and water masses on a single month-long expedition, sampling more densely along transects and deeper through the water column than possible on a global survey.

We undertook a high-resolution metagenomic survey of the Red Sea, conducting a multivariate community analysis of covariation between environmental parameters and metagenome-derived taxonomic and functional metrics. We followed three main lines of questioning. First, how well can both taxonomic and functional microbial diversity be explained by environmental parameters, and which environmental parameters explain the most variation? Sunagawa et al. (2015) showed in a recent global ocean survey that temperature could explain more variation in taxonomic abundance than any other parameter. At smaller spatial scales and narrower temperature ranges, does temperature still have the most explanatory power? Which parameters can best explain residual variation? Second, are patterns of environmental covariation conserved across co-occurring water masses? Sampling the GAIW allowed us to determine whether this cooccurring water mass follows the same organizational principles (covariation with environmental parameters) as the native Red Sea water mass, across different taxonomic and functional metrics. Third, how are microbes functionally adapted along environmental gradients of irradiance, temperature, salinity, and nutrients, including extremes in these parameters? Do marine communities exhibit fine-scale genomic adapation to environmental parameters as has been observed between separate oceans? Our dataset has allowed us to address these questions, and supporting online resources will make the processed data avaible to the wider community for further investigations.

## Materials and Methods

### Oceanographic Sampling

Samples were collected aboard the R/V *Aegaeo* on Leg 1 of the 2011 KAUST Red Sea Expedition, 15 September-11 October 2011. At eight stations, 20 L seawater was collected from each of depths 10, 25, 50, 100, 200, and 500 m; in two cases (Stations 12 and 34) where the seafloor was shallower than 500 m, the deepest sample was taken at the seafloor. Water was collected in 10-L Niskin bottles (i.e., two Niskin bottles per depth), attached to a CTD rosette. Back on deck, the seawater was filtered through a series of three 293-mm mixed cellulose esters filters (Millipore, Billerica, MA) of pore sizes 5.0 μm, 1.2 μm, and 0.1 μm. Filters were placed in sealed plastic bags and frozen at −20 °C. Station properties (location, depth of mixed layer, chlorophyll maximum, and oxygen minimum) are described in Table S1. Physical oceanographic measurements (pressure, temperature, conductivity, chlorophyll *a*, turbidity, and dissolved oxygen) were collected on a modified SeaBird 9/11+ rosette/CTD system, described in Supplementary Methods. Nutrient measurements (nitrate+nitrite, nitrite, ammonium, phosphate, and silicate) on the final 0.1-μm filtrate were carried out at the UCSB Marine Science Institute and the Woods Hole Oceanographic Institution (Supplementary Methods). Sample water properties are described in Table S2.

### DNA Extraction and Whole-Genome Shotgun (WGS) Sequencing of Community Metagenomes

Community DNA was extracted from the 0.1-μm filters (0.1-1.2 μm size fraction) using phenol-chloroform extraction, similar to Rusch et al. (2007) and Ngugi et al. (2012); the full protocol is described in Supplementary Methods. Yields of genomic DNA ranged from 200-1500 ng per sample. WGS libraries were made using the Nextera DNA Library Prep Kit (Illumina, San Diego, CA). Median insert size by sample ranged from 183-366 bp (Table S3). Libraries were sequenced using Illumina HiSeq 2000 paired-end (2 × 100 bp) sequencing, filling a total of three lanes (15 samples/lane). Sequence length after adapter removal was 93 bp, and 10 million reads (for each of reads 1 and 2) per sample were generated (Table S3). Reads were quality filtered and trimmed using PRINSEQ (Schmieder and Edwards, 2011) with parameters given in Table S4, and final read counts and metagenome sizes are given in Table S3. Although exact duplicates and reverse-complement exact duplicates were removed, we tested the effect of leaving in these duplicates, and it increased the number of reads retained by only 0.1–0.2%. Raw fastq files have been submitted to the NCBI BioSample database with accession numbers PRJNA289734 (BioProject) and SRR2102994-SRR2103038 (SRA). All analyses presented here were carried out on the quality filtered and trimmed reads. Both reads 1 and 2 were analyzed initially; however, unless otherwise indicated, only the results of read 1 are presented here because of a high degree of redundancy between results of reads 1 and 2. Genomic assemblies were built from each sample; these assemblies were used to calculate insert sizes of metagenomic libraries (Table S3) but provided limited value for quantitation of taxa and gene ortholog groups. The assemblies did, however, yield contigs belonging to uncharacterized clades, which are the subject of a separate study (Haroon et al., in review).

### Calculation of Metagenomic ‘Response Variables’ from Metagenomic Reads

Data tables of merged environmental metadata and response variables are provided in Supplementary Information. Scripts used in the preparation of this manuscript are available at https://github.com/cuttlefishh/papers in the directory red-sea-spatial-series.

*Taxonomic composition*. The 45 metagenomes were analyzed at the read level for the relative abundance of taxonomic groups using CLARK. CLARK (full mode) (Ounit et al., 2015) and CLARK-S (spaced mode) (Ounit and Lonardi, 2015) were used to classify paired metagenomic reads at species and genus level, respectively, based on a k-mer approach against the NCBI RefSeq database (Release 74). CLARK was run using default parameters but with the-highconfidence option, which reports only results with high confidence (assignments with confidence score >= 0.75 and gamma value >= 0.03), as suggested by the developers. For both species-level and genus-level CLARK results, the column Proportion_All(%) (relative normalized abundance such that each sample sums to 100%) was exported and merged with sample metadata (environmental parameters) using the Python package Pandas. Hierarchically-clustered heatmaps were generated using MetaPhlAn2 utilities (Truong et al., 2015).

In order to specifically capture the diversity within the *Pelagibacter* and *Prochlorococcus* groups in the Red Sea, we used GraftM (https://github.com/geronimp/graftM), which classifies reads based on HMM profiles in concert with a reference phlogeny. HMM profiles of *Pelagibacter* 16S rRNA gene and *Prochlorococcus rpoCl* were generated from forward reads using HMMer v3.1b1 (Eddy, 2011). Reference phylogenies were constructed using MEGA6 (Tamura et al., 2013) from ClustalW alignments (Larkin et al., 2007) of publicly available *Pelagibacter* 16S rRNA gene sequences (Luo et al., 2015) and *Prochlorococcus rpoCl* (DNA-directed RNA polymerase subunit gamma) genes (Shibl et al., in review). Phylogenies were estimated by maximum-likelihood using the GTR+I+G model of nucleotide evolution, chosen with the Perl script ProteinModelSelection.pl that comes with RAxML (Stamatakis 2014). GraftM was run with default parameters based on the the built GraftM packages, which are available here: https://github.com/fauziharoon/graftm-packages. Counts were fourth-root transformed.

*Gene ortholog group and pathway relative abundance*. The 45 metagenomes were analyzed for the relative abundance of gene ortholog groups (KEGG orthologs or KOs) and biochemical pathways (KEGG pathways) using HUMAnN v0.99 (Abubucker et al., 2012) with KEGG release 66.0. First, because the focus of this study was prokaryote genomes, and to increase search speed, reads were recruited to only the prokaryotic fraction of the KEGG genome database, containing all (as of the KEGG release) 1377 prokaryotic genomes (proteomes translated from open reading frames) using a translated search with USEARCH v7.0.1001 (Edgar, 2010) with options-ublast, -accel 0.8, and -evalue 1e-5. The fraction of reads mapped to the KEGG genome database averaged 26.2% (range 16.2–42.5%) across 45 samples (Table S6). Using these results, HUMAnN was run in both standard mode (all taxa merged) and in “per-organism” mode (option: c_fOrg = True). KO counts and KEGG pathway counts were normalized to counts per million (CPM) counts, i.e., the number of reads mapped to the KO (or pathway) divided by the sum of all reads mapped in that sample times 1e6, such that all values for a given sample sum to 1 million. Note that KO counts were not normalized to gene size (e.g., average length of each KO in KEGG) because this was unnecessary: comparisons of KO relative abundances were to environmental parameters and not to each other, and the multivariate community models used are insensitive to absolute magnitudes.

### Statistical Analyses

We utilized multivariate statistical techniques to relate an array of environmental parameters to metagenomic response variables: taxon relative abundance, KO relative abundance, and pathway relative abundance. All analyses were completed using R v3.1.1 (www.r-project.org) and PERMANOVA+ (Anderson et al., 2008).

*Exploratory analyses*. Pearson correlations between pairwise combinations of environmental parameters were calculated and displayed as a heat map. Similarity profile analysis (SIMPROF) was used to identify significant groupings within the KO relative abundance response matrix using the *clustsig* package (http://cran.r-project.org/web/packages/clustsig/index.html). Partitioning around medoids (PAM) was used to partition the KOs by relative abundance using the *cluster* package v1.15.2 (Kaufman and Rousseeuw, 2005) with Kullback-Leibler distances (Kullback and Leibler, 1951); 12 clusters were chosen based on minimization of the gap statistic.

*Explaining variability using environmental parameters* To quantify the spatial variation (both horizontally and vertically) in the response variable matrices explained by the co-occurring gradients in our environmental parameters, we used a multivariate distance-based linear model (DistLM) (McArdle and Anderson, 2001). Eight environmental parameters were considered: temperature, salinity, dissolved oxygen, chlorophyll, turbidity, nitrate, phosphate, and silicate. These parameters were normalized and fitted in a conditional manner to each response variable matrix using step-wise selection and 9999 permutations of the residuals under a reduced model. Model selection was based on Akaike’s information criterion with a second-order bias correction applied (AICc) (Hurvich and Tsai, 1989). The best-fit model (the one that balanced performance with parsimony) was then visualized using distance-based redundancy analysis (dbRDA) (McArdle and Anderson, 2001) in order to identify the directionality of the correlations between the response variable matrix and the environmental parameters. Variation explained by all parameters combined was calculated by forcing all parameters to be included in the final model.

*Visualization of metagenome-environment relationships*. Pairwise relationships between environmental parameters and KO relative abundance plus other metagenomic response variables were visualized using scatter plots, available using Bokeh-based HTML files in Supplementary Information. Environmental parameters and metagenomic response variables were visualized in the 3D volume of the Red Sea using the ili Toolbox, also available in Supplementary Information. KOs having strong correlations with environmental parameters were visualized with canonical correspondence analysis (CCA) using the *vegan* 2.3-1 package, implemented according to Legendre and Legendre (2012). For clarity, only KOs with abundance in the top half and variance in the top tenth of all KOs were visualized by CCA (Supplementary Methods).

## Results & Discussion

### Overview of Red Sea Metagenomic Dataset and Analysis

To measure covariation of microbial diversity with oceanic gradients, we sampled a north-south transect of the Red Sea at eight stations (Table S1), sampling six depths from the surface to 500 m (Figure 1A), totaling 45 samples. Concurrent with microbial sampling we measured temperature, salinity, dissolved oxygen, chlorophyll *a*, turbidity, nitrate, phosphate, and silicate (values in Table S2, covariance matrix in Figure 2). The microbial size fraction (0.1–1.2 μm) was sequenced at 10M reads per sample with 93-bp paired reads (Table S3). From the metagenomic reads, we calculated five metagenomic response variables: genus-level taxon relative abundance, species-level taxon relative abundance, gene ortholog group (KEGG Orthology or KO) relative abundance, KEGG pathway coverage, and KEGG pathway relative abundance. Of the 1738 taxa, 5775 KOs, and 162 pathways detected in the metagenomes, many exhibited ecologically meaningful correlations with environmental parameters. As an example, the inverse relationship between phosphate concentration and relative abundance of phosphate-acquisition gene *pstS* (K02040) is shown in Figure 1. Samples generally grouped by depth, as indicated by hierarchical clustering of samples based on all taxa (Figure 3) and KOs (Figure S1), and by abundance patterns of individual taxa and KOs (Figures 1B and Figures 5 and Supplementary Information).

**Figure 1.**
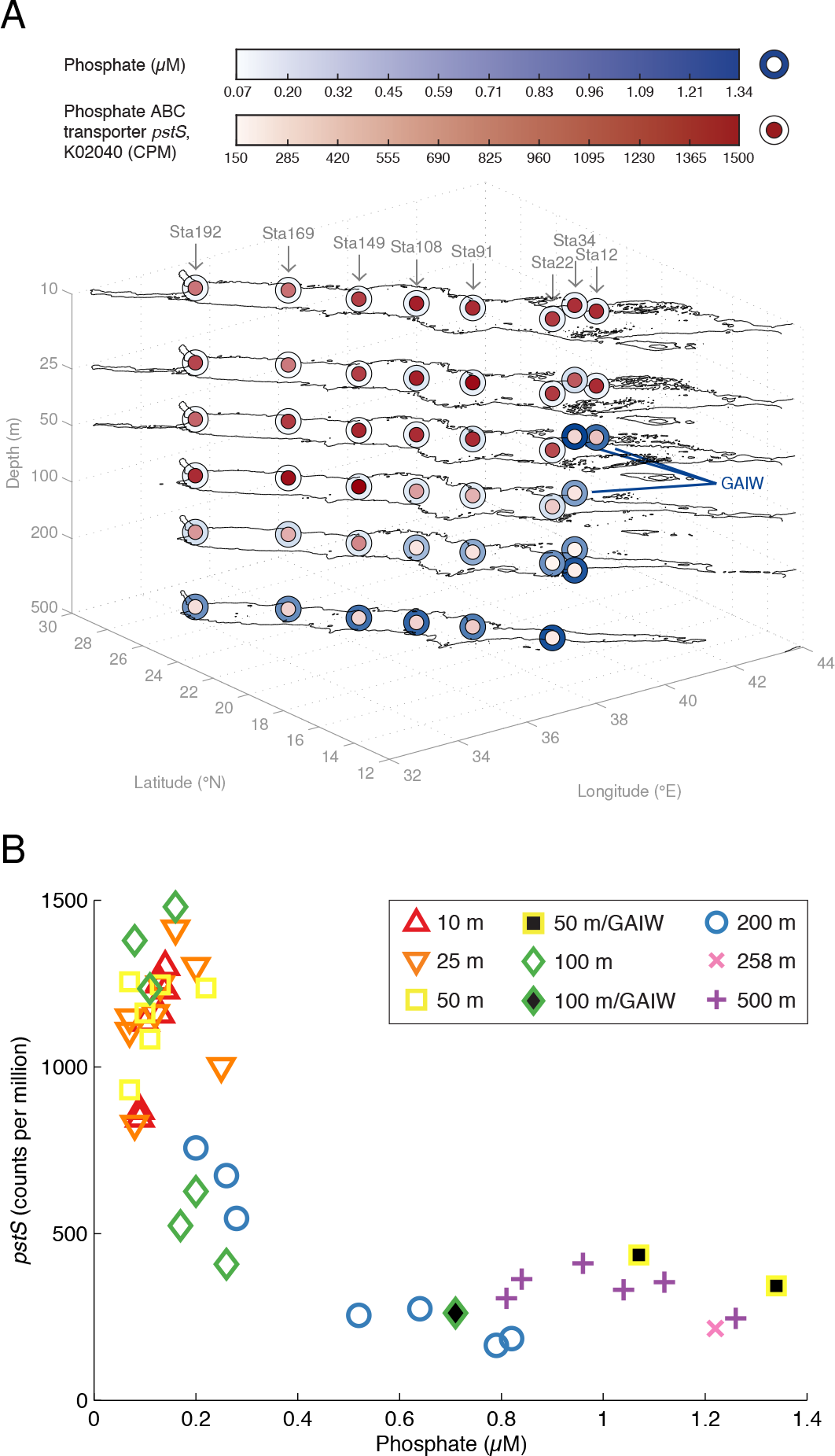
Covariation of gene ortholog group abundance and environmental parameters in the water column. (A) 3D contour map of the Red Sea, with outlines (isobaths) showing boundaries of the Red Sea at sampling depths, and samples colored by phosphate concentration (outer circle) and relative abundance of gene ortholog group (KO) for phosphate ABC transporter*pstS* (inner circle). (B) Scatter plot of KO relative abundance versus phosphate concentration. Samples taken within the foreign water mass Gulf of Aden Intermediate Water (GAIW) are indicated. KO relative abundance is given in units of counts per million (CPM) of total KO counts in each sample (i.e., all KOs sum to 1 million in each sample).

**Figure 2.**
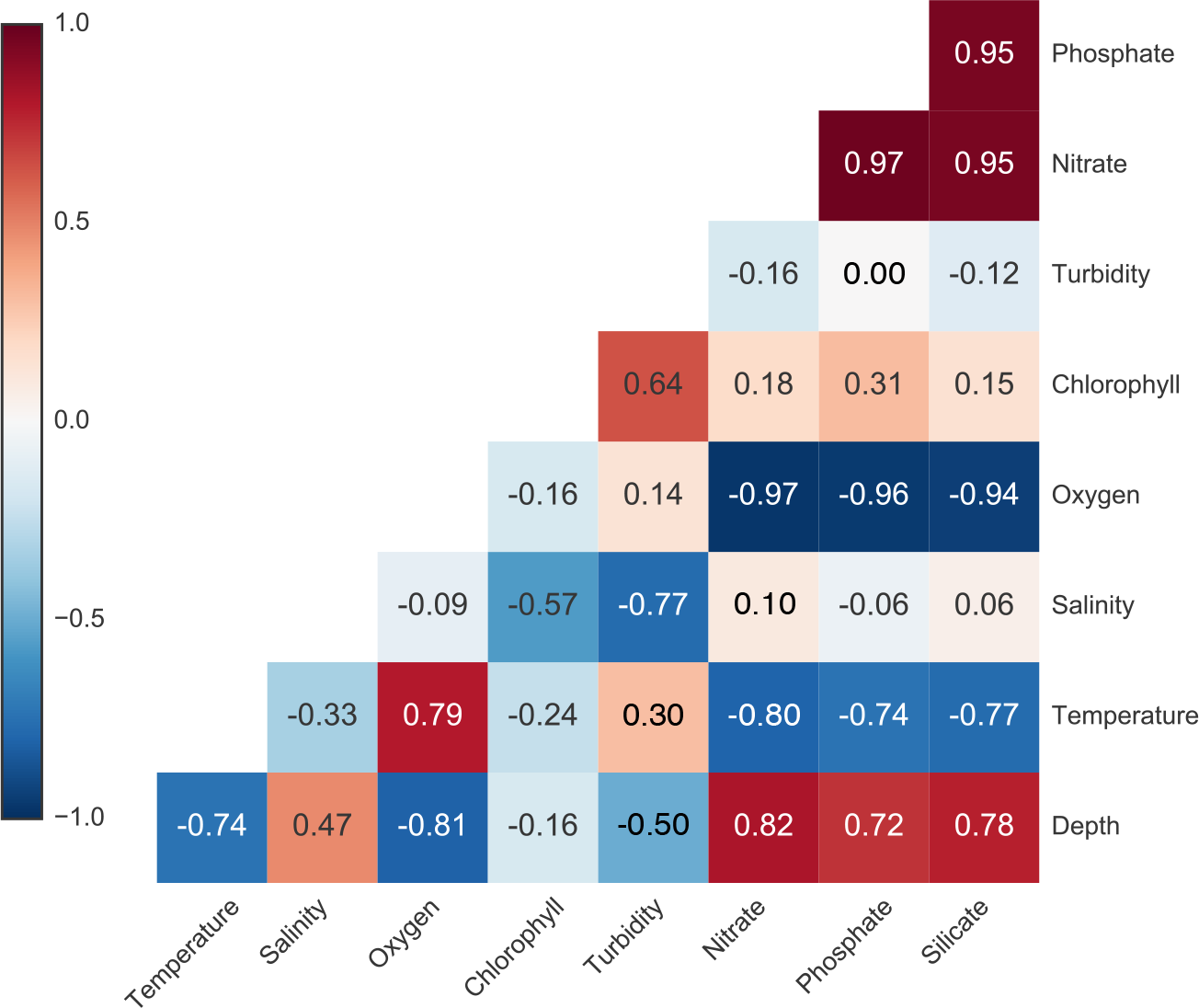
Pearson correlations between environmental parameters shown as a colored covariance matrix. A Pearson’s *r* value of 1 (red) indicates a total positive correlation, a value of −1 (blue) indicates a total negative correlation, and a value of 0 (white) indicates no correlation.

**Figure 3.**
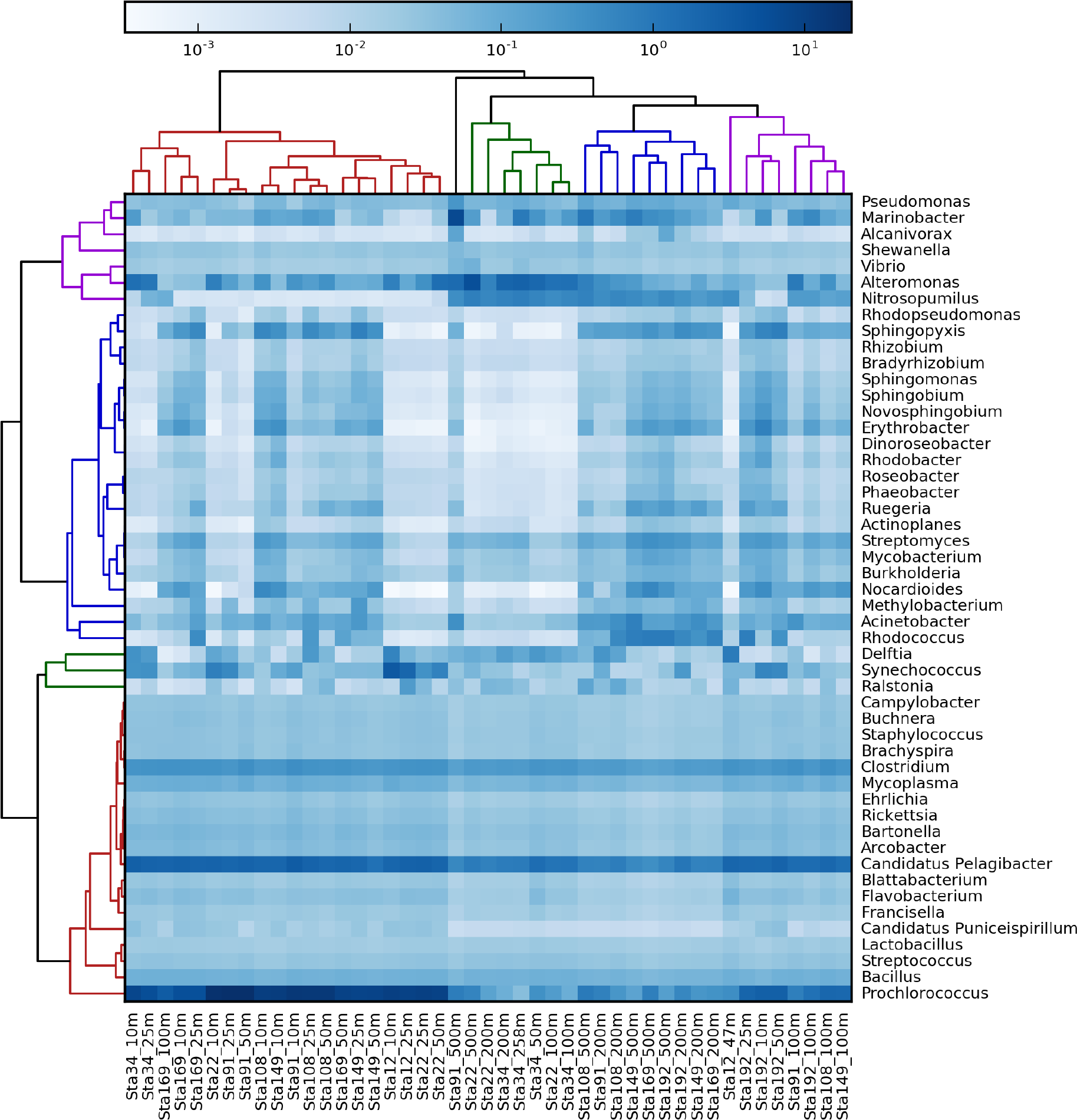
Relative abundance of genera across metagenomes displayed as a hierarchically-clustered heatmap, with clustering of samples by Bray-Curtis distance (top dendrogram) and clustering of taxa by correlation between samples (left dendrogram), with branch colors indicating major clusters. GAIW sample labels are colored red. The top 50 most abundant genera are shown. Relative abundances of all 683 genera detected for each sample sum to 100. Genus-level taxonomy was calculated based on k-mer frequency in comparison with the NCBI RefSeq database (methods).

The acquired set of metagenomic response variables and environmental parameters allowed us to assess the predictive power of environmental parameters at multiple levels of microbial genotype. We tested how much variation in genus-level taxonomy, KO relative abundance, and pathway relative abundance could be explained using a small number of environmental parameters. Distance-based multivariate linear models (DistLM) and redundancy analysis were used, balancing parsimony and performance (using AICc) to derive an optimal model for explaining variation in each response variable (Figure 4). We acknowledge that the analyses presented here, by necessity, are constrained by the databases available for assigning taxonomy and KOs and the available mappings of KOs to pathways.

**Figure 4.**
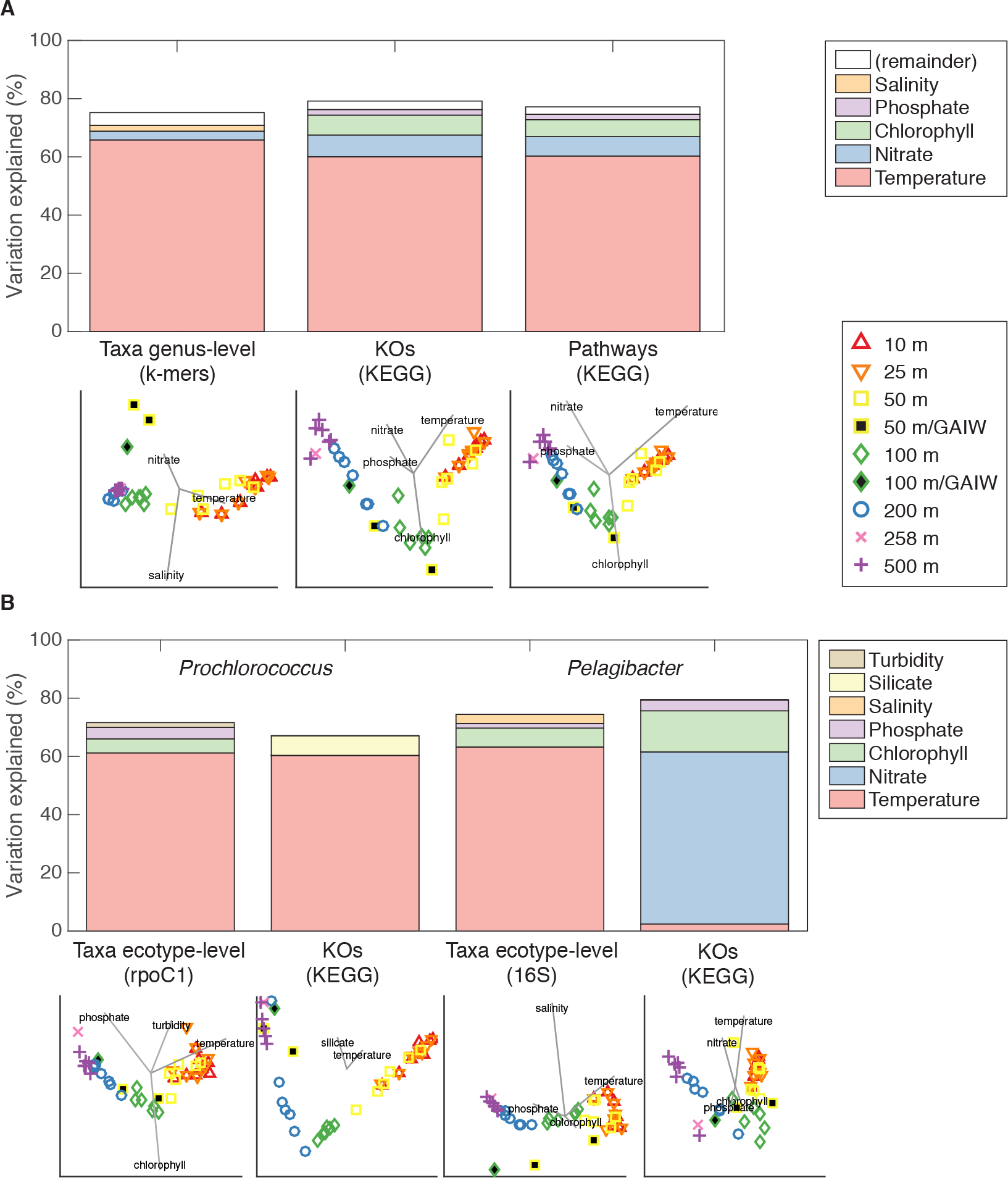
Maximization of linear relations between environmental parameters and metagenomic response variables using a distance-based multivariate linear model and distance-based redundancy analysis for (A) the whole data set and (B) genera *Prochlorococcus* and *Pelagibacter* (see methods). Percent variation explained by each parameter is shown as a bar graph. The optimal model using AICc to balance performance and parsimony is shown for both (A) and (B); also shown for (A) is the remainder of variation explained by other environmental parameters unused in the optimal model. The dbRDA ordination of the optimal model is shown along dbRDA axes 1 and 2, with stations colored by depth and water mass (GAIW in black).

### Variation in Metagenomic Diversity Metrics Explained by Environmental Parameters

We first asked which environmental parameters explained the most variation in both taxonomic and functional diversity metrics, and we looked for differences in total variation explained. Environmental parameters explained similar amounts of variation in the various metrics used (Figure 4A). Total variation explained using all available environmental parameters was only marginally higher for KO relative abundance (79.0%) than for pathway relative abundance (77.0%) and genus-level taxon relative abundance (75.1%). Variation explained was similar even at greater phylogenetic resolution within two important marine microbial groups, the autotroph *Prochlorococcus* and the heterotroph *Pelagibacter* (SAR11 clade), which are the two most abundant genera across our dataset (Figure 3). At ecotype-level taxonomy *(Prochlorococcus* “ecotypes” and *Pelagibacter* “subclades”) and genus-level KO abundance, the percent variation explained was similar to the community as a whole (Figure 4B).

Overall, environmental parameters explained more variation in our dataset than in other microbial ecosystems where this has been tested. For example, in a similarly sized dataset on reef-associated microbes, the best parameter explained only 15% of taxonomic variation and 18% of metabolic variation (Kelly et al., 2014). Variations in water column microbial communities appear easier to predict. In an English Channel time-series, day length explained over 65% of variance in taxonomic diversity (Gilbert et al., 2011). The better performance of water column data could be because the open ocean is not patchy but well mixed and stably stratified by depth into layers, especially in the Red Sea in late summer. Relative to open-water samples, the increased complexity of the response matrix in reef-associated samples resulting from microhabitats and higher diversity likely reduces model performance.

Temperature explained the most variation in each of the response variables; this was followed in each case by nitrate (second) and then chlorophyll (third) for the functional response variables and salinity (third) for genus-level taxonomy (Figures 4A and Figures S2). Although nitrate and phosphate (*r*=0.97) and silicate (*r*=0.95 with nitrate and phosphate) were very highly correlated (Figure 2), nitrate was consistently ranked higher (explaining more variation) than phosphate in the optimal model, and silicate was not implicated (Figure 4A). Across the whole dataset, temperature explained more variation than oxygen in every response variable. Although temperatue and oxygen were correlated (*r*=0.79), oxygen was never part of the optimal model. Temperature has been identified as a key predictor of microbial diversity in the ocean by other studies (Johnson et al., 2006; Sunagawa et al., 2015). Specifically, Sunagawa et al. (2015) showed that temperature is a better predictor of taxonomic composition than is oxygen. Here we show that the same is true for gene functional composition (KOs): the absence of oxygen in any optimal model suggests that temperature is a stronger predictor (and possibly also driver) of microbial diversity than oxygen.

Nitrate (measured as nitrate+nitrite) and phosphate both formed part of the optimal model for each functional response variable, with nitrate always explaining slightly more variation than phosphate. This finding hints at the relative selective pressures these two key nutrients exert. The idea that limitation of a given nutrient leads to the gain of genes for uptake and assimilation of that nutrient is supported by numerous studies (Coleman and Chisholm, 2010; Kelly et al., 2013; Thompson et al., 2013). Here we extend that idea to the quantitative explanatory power of the nutrient’s concentration for predicting KO relative abundance. The predictive power of nitrate relative to phosphate in our genetic results may indicate that nitrogen (N) is relatively more limiting than phosphorus (P) in the Red Sea. Limited data exist on this topic, but N:P ratios of 0.3–5 (well below the Redfield ratio of 16, the atomic ratio of N to P in phytoplankton (Redfield, 1958)) in the Gulf of Aqaba (Lindell et al., 2005) and a high frequency of N-acquisition genes in a Red Sea surface metagenome relative to the Atlantic ocean (Thompson et al., 2013) suggest N limitation; however, in the northern Gulf of Aqaba, a P-stress response and lack of N-stress *ntcA* response in Red Sea cyanobacteria supports the opposite conclusion (Post, 2005). Nevertheless, our own nutrient measurements from this cruise show that the N:P ratio (calculated here as the ratio of nitrate+nitrite to phosphate) in surface waters was 2, whereas a prototypical ratio of 16 was observed in deeper waters (Figure S3), possibly due to remineralization of N from phytoplankton at depth. Regarding nitrate, it is interesting that for *Pelagibacter* KO relative abundance, nitrate (59.1%) not temperature explained the most variation. N limitation has strong effects on the transcriptional response of *Pelagibacter* in culture, with genes for assimilation of organic sources of N up-regulated under N stress (Smith et al., 2013. However, none of the differentially expressed genes identified by Smith et al. (2013) covaried strongly with nitrate in our dataset (reads from corresponding KOs assigned to *Pelagibacter*). Thus, the nature of a potential selective force of this putative N limitation on *Pelagibacter* gene content remains a mystery.

Chlorophyll has a non-monotonic relationship with depth, unlike the other environmental parameters analyzed here, which are either low at the surface and high at depth (salinity, phosphate, nitrate, silicate) or high at the surface and low at depth (temperature, oxygen, turbidity). Chlorophyll peaks below the surface mixed layer at the deep chlorophyll maximum (100 m in the Red Sea, Table S1), due to a confluence of sunlight from above, nutrients from below, and the tendency of deeper phytoplankton to possess higher chlorophyll per cell. Because chlorophyll is effectively orthogonal to other environmental parameters, it should not be unexpected that it has significant explanatory power, and that chlorophyll and temperature (a key depth-dependent parameter) together could explain much of the genetic variation.

### Comparison with a Foreign Water Mass

We next asked whether the ability to predict metagenomic response variation from environmental parameters was sufficiently robust to extend to alternate water masses. Fortuitously, the Red Sea experiences a water influx each summer from the Indian Ocean, called the Gulf of Aden Intermediate Water (GAIW), which was captured in three of our samples. The GAIW brings cooler, less saline, oxygen-rich, nutrient-rich water from the Gulf of Aden (Churchill et al., 2014). The three GAIW samples were clearly distinct from their neighboring samples in the T-S profile (Figure S4A) and Red Sea water column (Figure S4B). The properties of GAIW samples resembled those of deeper samples in the native Red Sea water mass; the GAIW samples, which were from 50–100 m depth, had markedly different environmental parameters from non-GAIW samples from 50–100 m. We were curious if our multivariate community models would be able to highlight any differences between response variables in model performance across different water masses.

Considering the distance-based redundancy analysis (Figures 4A and Figures S2), all of the functional metrics placed the GAIW samples amidst the native Red Sea samples, though clustering with deeper samples, owing to the lower temperature and higher nutrients of the GAIW samples. The taxonomic metrics, however, placed the GAIW samples either far apart from the other samples (genus-level) or with much deeper samples than expected even based on physicochemical properties (species-level), driven by the high nitrate and low temperature and salinity of the GAIW samples relative to the non-GAIW samples(Figure 4A). These results suggest that environmental covariation patterns of taxonomy are less conserved across water masses (i.e., different combinations of environmental parameters) than are environmental covariation patterns of functional metrics.

Supporting the idea that functional covariation with environmental parameters is conserved across different water masses, we note anecdotally that for most of the individual environment–KO relationships examined below (Figure 5), GAIW samples followed a similar pattern to the non-GAIW majority. One notable exception was salinity, with the salinity of GAIW much lower than anything in the native Red Sea water mass and the covariation of KOs with salinity very different for GAIW samples compared to non-GAIW samples.

**Figure 5.**
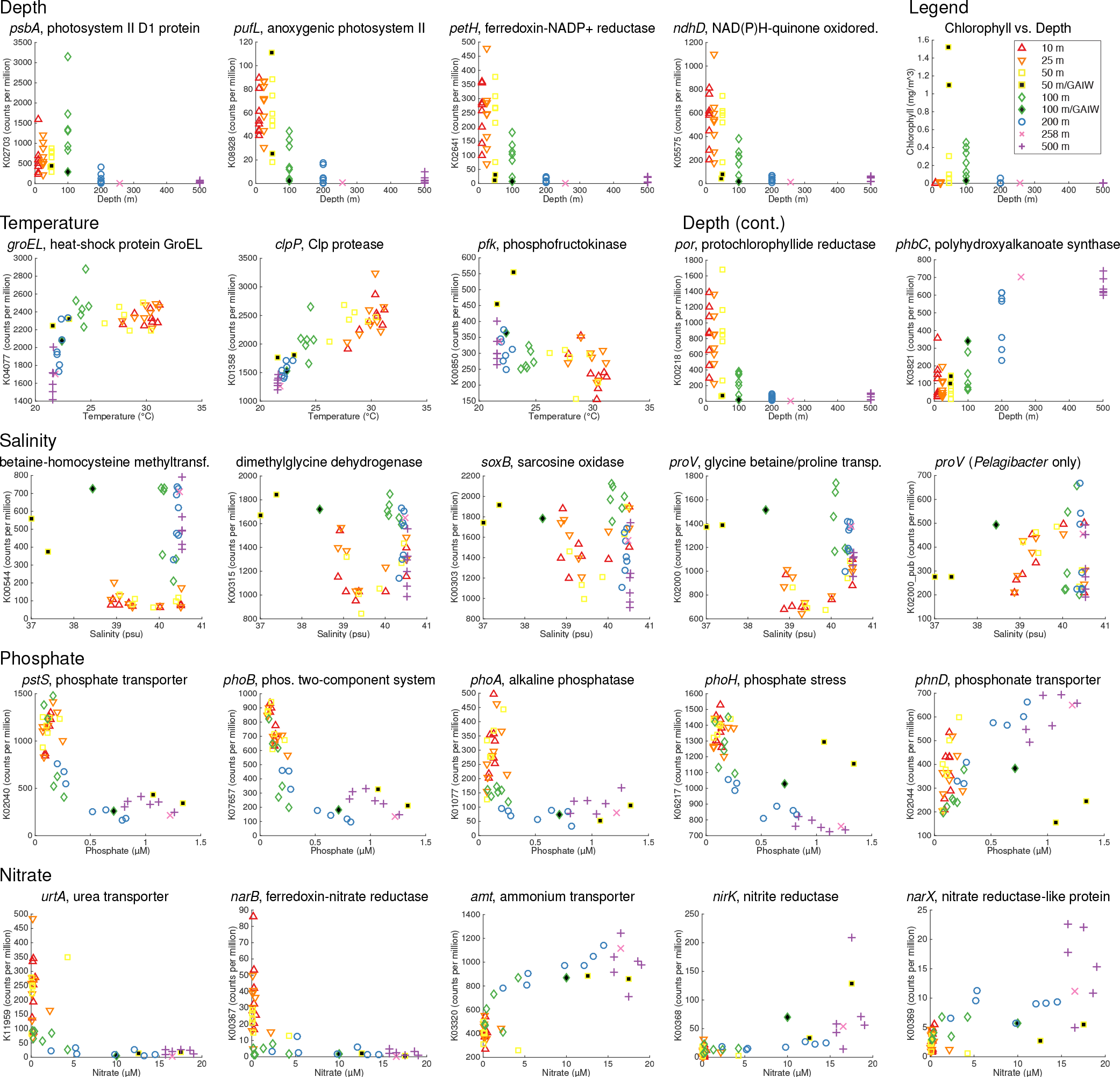
Covariation of select KOs with environmental parameters. KO relative abundance is given in units of counts per million of total KO counts in each sample (i.e., all KOs sum to 1 million in each sample).

### Environmental Covariation Patterns of Individual KOs

We finally turned our attention to the covariation patterns of individual KOs, which partition along the three-dimensional water column in ecologically meaningful ways. Which KOs have the strongest covariation with environmental parameters? Can previously observed patterns between oceans also be observed along gradients within a single sea? Which KOs are implicated in the adaptive response of microbes to the low nutrients and high irradiance, temperature, and salinity of the Red Sea?

We used canonical correspondence analysis (CCA) to identify and visualize correlations between KOs and environmental parameters, with KOs organized by metabolic pathway (Figure 6 and Table S9). We note that all KOs were included in the distance-based linear model above, whereas a subset of the most differentially represented and abundant KOs are shown in the CCA (methods); most of the KOs discussed below are visualized in Figure 6. Additionally, KOs were ranked by total abundance across all samples (Table S7), and KO abundance patterns were clustered using partitioning around medoids (PAM) into 12 clusters (Table S8).

**Figure 6.**
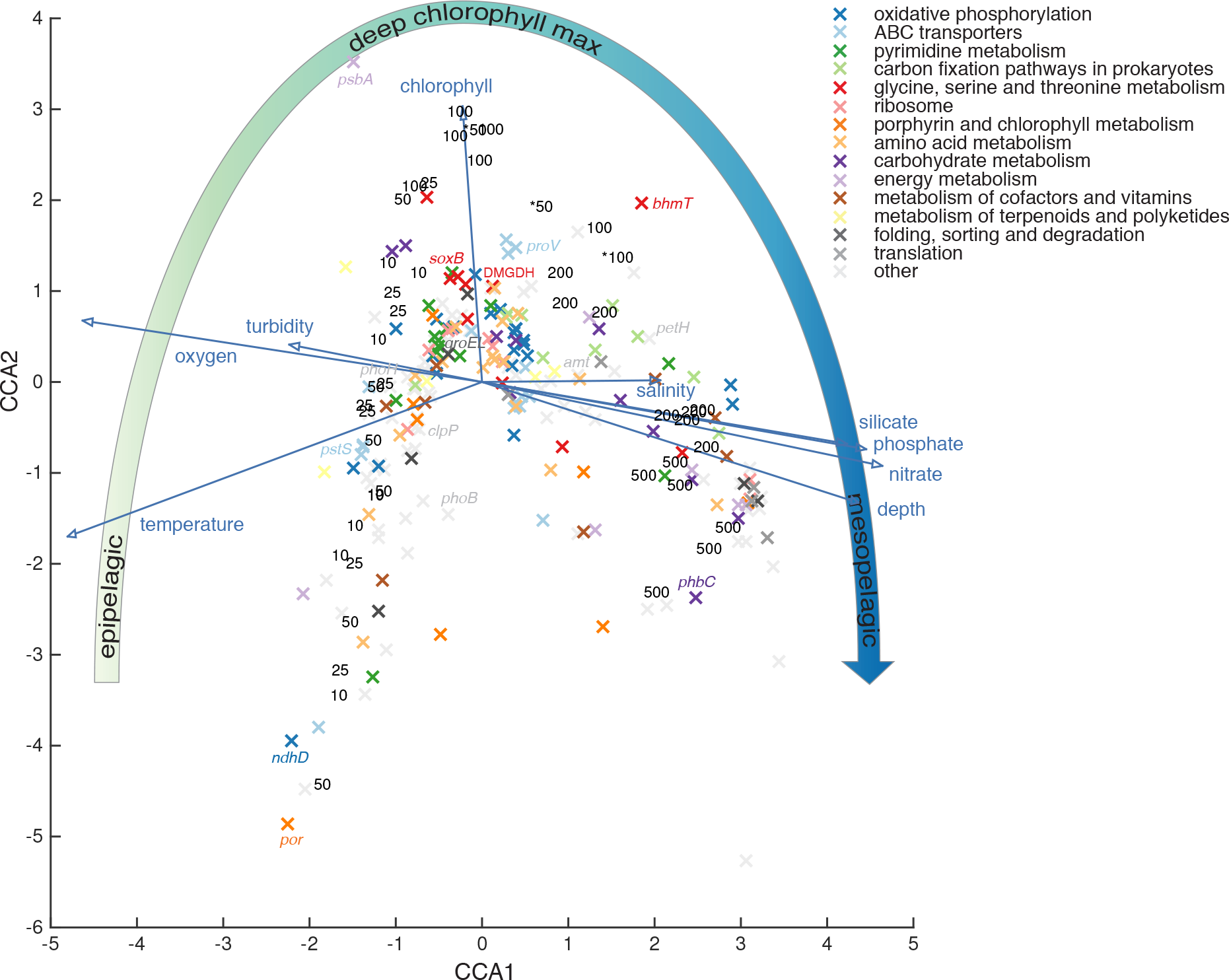
Canonical correspondence analysis of KO relative abundance with environmental parameters. Samples are shown as black numerals indicating depth in meters (GAIW samples marked with asterisk), environmental parameters as dark blue arrows, and KOs colored by KEGG pathway. For clarity, only KOs were displayed that were found in all samples, with a total count of at least one per thousand counts over all samples, and variance in the top 10% (see methods). The large arrow indicates the trend of sample position from surface (epipelagic), to deep chlorophyll maximum, to deep (mesopelagic).

Nutrient acquisition and energy metabolism contributed most of the functional covariation with environmental parameters (Figure 6). The patterns documented here were not exclusively depth-dependent but also captured subtle covariation with gradients along isobaths. The full set of environmental parameters and metagenomic response variables can be visualized interactively in the 3D volume of the Red Sea using web-based tools with files in Supplementary Information. Visualization examples showing the temperature-salinity (T–S) profile and temperature in 3D are provided in Figure S4.

Depth is a spatial parameter that is not ‘felt’ by microorganisms, except as it relates to pressure, but nevertheless structures virtually all environmental parameters in the water column. Light attenuates with depth, and thus photosynthesis is mostly confined to the upper 200 m of the water column. As expected, KOs for photosynthesis were most abundant in shallow waters and gradually less abundant in deeper waters. This was true for both oxygenic (*psbA/*K02703) and anoxygenic (*pufL/*K08928) photosynthesis, photosynthetic electron transport (*petH/*K02641, *ndhD/*K05575), and pigment biosynthesis (*por/*K00218) (Figure 5). Some heterotrophic bacteria accumulate carbon-rich polymers called polyhydroxyalkanoates when organic carbon is readily available but growth is limited by nutrients (Stubbe et al., 2005). We observed that polyhydroxyalkanoate synthase (*phbC/*K03821) was more abundant in mesopelagic samples than epipelagic samples, consistent with relatively more heterotrophy than phototrophy at depth.

Temperature covaries with depth (warmer at surface, colder at depth), but in the Red Sea southern surface waters are warmer than in the north, and the GAIW is cooler than surrounding depths in the native Red Sea water mass. We observed that KOs for chaperonins, including heat-shock proteins GroEL/ES (K04077, K04078), and proteases, including Clp protease (*clpP/*K01358), had greater relative abundance in warm (24–32°C) samples than cooler (21–23°C) samples (Figure 5). Both GroEL/ES (Zeilstra-Ryalls et al., 1991) and Clp protease (Zybailov et al., 2009) have important roles in protein folding, which is sensitive to high temperature. The increase in *groEL* relative abundance leveled off above 23 °C, whereas the increase in *clpP* relative abundance increased along the full temperature range from 21 to 32 °C. In an opposite trend, glycolysis was relatively more abundant in colder samples than warmer samples, as exemplified by phosphofructokinase (*pfk/*K00850). This is likely related to the relative increase in heterotrophy at depth, as deeper waters tend to be cooler. The most cold, eutrophic samples from the GAIW have the highest relative abundance of *pfk* by far, indicating relatively more heterotrophy in this foreign water mass.

Salinity in the Red Sea is higher at depth and in northern surface waters and lower in southern surface waters and the GAIW. Saline-rich waters of the the Mediterranean and Red Seas were previously shown to have high relative abundance of genes for degradation of osmolytes, in particular recruiting to *Pelagibacter* (Thompson et al., 2013). We put forth a hypothesis that high salinity leads to high production of osmolytes by algae and other organisms, a valuable organic carbon and nutrient source for *Pelagibacter* (Sun et al., 2011), and therefore there is selective pressure to encode osmolyte-degrading enzymes (Thompson et al., 2013). Across the 45 Red Sea metagenomes, KOs for glycine betaine (GBT) transport and degradation (Sun et al., 2011) -glycine betaine/proline transporter (*proV/*K02000), betaine-homocysteine S-methyltransferase (*bhmT/*K00544), dimethylglycine dehydrogenase (DMGDH/K00315), and sarcosine oxidase (*soxB/*K00303) -were correlated with high chlorophyll and with high or moderate salinity (Figure 6). The shape of covariation of these four KOs was not as clearly dependent on salinity as we expected (Figure 5). As suggested by the CCA plot (Figure 6), both salinity and chlorophyll help explain the relative abundance of GBT transport and degradation KOs. The legend at top-right of Figure 5 indicates chlorophyll *a* fluorescence of the samples as a function of depth. Among the GBT-utilization KOs, samples with either high chlorophyll (green and yellow-black) or high salinity (blue and purple) tended to have the highest abundance. Thus this multifaceted trend is completely consistent with the hypothesis of phototroph (chlorophyll) production of osmolytes in high-salinity waters as a source of reduced carbon and nutrients for heterotrophic bacteria. Regarding the phototrophs responsible for producing GBT, we note that while *Prochlorococcus* are thought to use glucosylglycerate and sucrose as their main osmolytes, some low-light *Prochlorococcus* strains and *Synechococcus* strains are thought to accumulate GBT as well (Scanlan et al., 2009), and these low-light strains are more abundant in the high-chlorophyll samples. Interestingly, the KO pattern with reads assigned to *Pelagibacter* specifically (e.g., *proV/*K02000) – one- to two-thirds of the recruited reads for these salinity-related KOs — were similar to the overall KO pattern but more dependent on salinity than chlorophyll (Figure 5).

Phosphate and nitrate are both low in Red Sea surface waters but higher at depth and in the GAIW (for example, phosphate shown in Figure 1A). Several studies have shown that genes for nutrient acquisition are enriched in waters limited for those nutrients, e.g., phosphate acquisition in the low-phosphate Mediterranean and Sargasso Seas (Coleman and Chisholm, 2010; Kelly et al., 2013; Thompson et al., 2013). Across the gradients of the Red Sea, numerous KOs for nutrient transport and assimilation were differentially distributed between nutrient-poor (surface and non-GAIW) and nutrient-rich (deep and GAIW) samples. Although depth was a major factor underlying the covariation observed here, we also detected more subtle differences along gradients within isobaths, as well as more striking differences between GAIW and non-GAIW samples at the same depth. This is to our knowledge the first demonstration of differential abundance patterns of nutrient-acquisition genes on such a small scale, not between disparate oceans but across environmental gradients within a single sea.

Phosphate-acquisition and phosphate-response KOs were enriched in low-phosphate samples (Figure 5), including phosphate ABC transporter (*pstS/*K02040), phosphate two-component system PhoBR (K07657, K07636), alkaline phosphatase (*phoA/*K01077), and phosphate stress-response protein PhoH (K06217). Trends were observed even within isothermal samples binned in two-degree increments, both for cooler isotherms with a wide range of phosphate concentrations, and for warmer isotherms with a narrow and low range of phosphate concentrations, e.g. *phoB/*K07657 (Figure S5). Phosphonate-acquisition genes, in an opposite pattern, were enriched in high-phosphate (and low-chlorophyll) samples, as exemplified by the phosphonate ABC transporter (*phnD/*K02044). Phosphonate utilization genes (*phn*) are abundant in proteobacteria such as *Pelagibacter* (Villarreal-Chiu et al., 2012), and are enriched in deeper waters of the Sargasso Sea (Martinez et al., 2010) and generally in low-P waters (Coleman and Chisholm, 2010; Feingersch et al., 2010). Although phosphonate-acquisition genes are found in some *Prochlorococcus* in the environment (Feingersch et al., 2012), genomes and transcriptional responses of cultured strains (Martiny et al., 2006) suggest that inorganic phosphate is the major P source for *Prochlorococcus*. Therefore, in addition to ecotype-level genome variability tuned to ambient concentrations of phosphate and phosphonate (Martiny et al., 2006), different distributions of phosphate- and phosphonate-acquisition genes along the water column are likely also due to genus-level differences in taxonomic composition (and therefore gene content) along the water column, for example, phosphate-utilizing *Prochlorococcus* in the epipelagic and phosphonate-utilizing *Pelagibacter* in the mesopelagic. Indeed, many of the low-nutrient-associated KOs such as phosphate and urea transporters had very similar abundance patterns to KOs typical of a phototrophic bacterium like *Prochlorococcus*: photosystems and photosynthetic electron transport, chlorophyll binding proteins, the Calvin cycle, and transport and chelation of metal cofactors essential for photosynthesis (PAM cluster 8, Table S8).

Nitrogen-acquisition KOs were differentially distributed with respect to nitrate concentration and, like with phosphorus, also followed one of two opposite patterns (Figure 5), which were also observed within isotherms (Figure S5). KOs for urea transport (*urtA/*K11959) and assimilatory ferredoxin-nitrate reductase (*narB/*K00367) were enriched in low-nitrate relative to high-nitrate samples. Conversely, KOs for ammonium transport (*amt/*K03320), nitrite reductase (*nirK/*K00368), and nitrate reductase-like protein (*narX/*K00369) were enriched in high-nitrate relative to low-nitrate samples, with the shift to high abundance occurring at 5 μM for *amt* and 15 μM for *nirK* and *narX*. Our measurements of N species besides nitrate were either below detection (nitrite) or unreliable (ammonium), but using global averages, nitrite and ammonium peak around the chlorophyll maximum and nitracline (where nitrate increases most rapidly) and then decrease through the deep epipelagic and mesopelagic (Gruber, 2008); urea is generally low and patchy through the water column (Remsen, 1971). Abundance patterns of several N-acquisition KOs thus appear to follow the “low nutrient-high KO” paradigm: nitrate reductase was abundant where nitrate was low (surface), and ammonium transport and nitrite reductase were abundant where ammonium and nitrite were low (mesopelagic). Urea transport, if it follows the same paradigm, indicates that urea in the surface of the Red Sea is also very low relative to at depth.

### Conclusions

We have analyzed a 3D array of marine metagenomes across environmental gradients in the Red Sea, showing that three-quarters of taxonomic and functional variation could be explained by temperature, nitrate, and chlorophyll. Covariation patterns with environmental parameters were largely conserved across water masses, notably more so for gene orthologs and pathways than for taxonomic groups. Individual patterns of KO covariation with environmental parameters revealed protein folding functions highly correlated with temperature, osmolyte degradation functions correlated with salinity and chlorophyll, and acquisition functions of nitrogen and phosphorus species anti-correlated with concentrations of their respective species. Subtle trends shown here across isobathic and isothermal gradients have hitherto been observed only between distant and disparate oceans. It is expected that this high-resolution marine metagenomic map of the Red Sea, accessible using interactive visualization tools, will serve as an important resource for marine microbiology and modeling

## Acknowledgments

We thank chief scientist Amy Bower, co-chief scientist Yasser Abualnaja, Leah Trafford, Dan McCorkle, and other scientists from the Woods Hole Oceanographic Institution, the captain and crew of the R/V *Aegaeo* and the Hellenic Center for Marine Research, and Red Sea Research Center Director James Luyten for their help on the 2011 KAUST (King Abdullah University of Science and Technology) Red Sea Expedition. Assistance with DNA extraction was provided by Matt Cahill, David Ngugi, and Francisco Acosta Espinosa. Bioinformatics assistance was provided by Mamoon Rashid and James Morton. Statistics assistance was provided by Mikyoung Jun, Myoungji Lee, Yoan Eynaud, and James Morton. We thank Jon Sanders, Jenan Kharbush, and Lihini Aluwihare for helpful comments on the manuscript. We also thank colleagues who suggested KOs hypothesized to have interesting ecological patterns: Paul Berube, Yue Guan, Laura Villanueva, Francisco Rodríguez-Valera, Nathan Ahlgren, Zhenfeng Liu, Francy Jiménez, and Ulrike Pfreundt. This work was funded in part by a postdoctoral fellowship to L.R.T. from the Saudi Basic Industries Corporation (SABIC).

## Author Contributions

L.R.T. planned the study, organized the cruise, collected samples, organized chemical analysis, curated physical and chemical data, extracted DNA, planned sequencing, processed sequence data, planned statistical analysis, made graphs and tables, and wrote the paper. G.J.W. planned and executed statistical analyses and wrote the paper. M.F.H. tested and ran taxonomic analyses and wrote the paper. A.S. collected samples and extracted DNA. P.L. ran metabolite prediction analysis. J.S. generated interactive visualizations. R.K. provided analytical input and wrote the paper. U.S. planned the study, organized the cruise and wrote the paper.

## Author Information

Sequence data have been submitted to the NCBI BioSample database with accession numbers PRJNA289734 (BioProject) and SRR2102994-SRR2103038 (SRA).

## Supplementary Methods

*Physical and chemical measurements of seawater*

Oceanographic measurements were collected on a modified SeaBird 9/11+ rosette/CTD system. Sensors on the CTD included those for pressure, temperature and conductivity (SBE9+ CTD), chlorophyll *a* and turbidity (WETLabs), and dissolved oxygen (SeaBird). Seawater samples were collected at nearly all stations for shipboard calibrations of salinity and oxygen CTD measurements. For nutrient measurements, filtrate from the final 0.1μm filters was collected and frozen at −20 °C, then analyzed by flow injection analysis at the UCSB Marine Science Institute (nitrate+nitrite, nitrite, ammonium, and phosphate) and the Woods Hole Oceanographic Institution (silicate). Ammonium was measured at less than 2 μM and nitrite at less than 0.7 μM for all samples; these data were determined to be insufficiently precise for statistical analyses because nitrite concentrations were near or below the detection limit and the ammonium measurements could not be done on fresh (unfrozen) samples.

### DNA Extraction

Filters were cut aseptically into small pieces and placed in tubes. Lysis buffer (0.1 M Tris-HCl, 0.1 M Na-EDTA, 0.1 M Na2H2PO4, 1.5 M NaCl, pH 8.0) was added to a volume of 15 mL. Three cycles of freeze-thaw (−80 °C, then 65 °C) were carried out. Lysozyme (2.5 mg/mL final conc.) and RNase A (2 mg/mL final conc.) was added, and the tubes were heated with rotation at 37 °C for 1 h. Next, proteinase K (0.2 mg/mL final conc.) and SDS (1% final conc.) was added, and the tubes were heated with rotation at 55 °C for 2 h. Lysate was extracted with an equal volume of phenol:chloroform:isoamyl alcohol (25:24:1), then an equal volume of chloroform:isoamyl alcohol (24:1). DNA was concentrated and washed with 10 mM TE (pH 8.0) using Amicon Ultra filters (10 kDa MWCO). Finally, DNA was ethanol precipitated for additional purity.

### Additional Statistical Exploratory Analyses

Testing for zonation patterns across depth gradients was carried out using a hypothesis-driven, constrained approach to test whether clear gradients existed in microbial diversity across depth (fixed factor), a prediction supported by numerous studies to date (DeLong et al., 2006; Ngugi and Stingl, 2012). We used a permutational multivariate analysis of variance (PERMANOVA) (Anderson, 2001), with subsequent pairwise comparisons, to formally test for differences in multivariate response variables across six a priori defined depth zones (10, 25, 50, 100, 200, 500 m). Analyses were based on Bray-Curtis similarity matrices, type III (partial) sums-of-squares, and unrestricted permutations of the raw data. The results of the PERMANOVA were visualized using a canonical analysis of principal coordinates (CAP) (Anderson and Willis, 2003), with allocation success (Anderson and Robinson, 2003) quantified across groups. Allocation success (expressed as a percentage) gave a measure of how distinct one depth group was relative to all other depth groups in the CAP multivariate space. Multidimensional scaling (MDS) was used to determine similarity between samples using the *vegan* package v2.3-1 (http://vegan.r-forge.r-project.org/).

### Alternate Taxonomy Composition Methods

In addition to CLARK (main methods), two other metagenomic taxonomy composition methods, described below, were tested. Based on the percent of reads mapping (Table S6), the percent taxonomic variation explained using environmental parameters, and an independent comparison of the methods (Lindgreen et al., 2016), we chose CLARK for determining taxonomic composition of each metagenomic library.

GraftM v0.6.3 (https://github.com/geronimp/graftM) employs a Hidden Markov Model (HMM) with pattern recognition to identify 16S rRNA gene sequences (https://www.orau.gov/gsp2015/abstracts/TysonG_02.pdf) and assign them to taxonomic groups based on tree placement using the SILVA database (Release 119, Quast et al. (2012)), to the most granular phylogenetic level possible using pplacer (Matsen et al., 2010). This non-BLAST approach allows more accurate classification regardless of incomplete databases (https://www.orau.gov/gsp2015/abstracts/TysonG_02.pdf). GraftM was then used with default parameters, using forward reads only. Total counts were presence/absence transformed or fourth root transformed.

Kraken (Wood and Salzberg, 2014) attempts to assign taxonomy to each read or read pair using k-mer analysis. Metagenomes were analyzed in ‘paired’ mode using the NCBI RefSeq database. Genus-level counts were normalized to the total number of paired reads per metagenome. Several key genera—*‘Candidatus* Pelagibacter ubique’, hereafter *Pelagibacter*, and groups of phages infecting *Prochlorococcus, Synechococcus*, and *Pelagibacter*—were manually curated to include taxa inadequately classified in NCBI Taxonomy. Total counts were normalized to total paired reads.

Kraken and GraftM were not selected because they mapped fewer sequence reads (avg. 8.8% for Kraken genus-level, 0.05% for GraftM genus-level, Table S6) and had lower percent variation explained. Further, a recent comparison of metagenomic taxonomy assignment tools found that CLARK performed best among tools tested in correlation between predicted and known relative abundances with a test data set (Lindgreen et al., 2016).

Total variation explained of species-level taxon relative abundance (Table S5) was similar (76.1%) to genus-level relative abundance (75.1%). Given this similarity, along with the two-fold greater number of reads mapped for genus-level (avg. 11.6%) than species-level (avg. 6.2%) assignment (Table S6) by the k-mer based CLARK software (Ounit et al., 2015) and the observed greater accuracy of genus-level taxonomic assignments, we chose to focus on genus-level taxonomy for analyses.

### CCA Visualization of Metagenome-Environment Relationships

The filtering described here was used only for selecting KOs to visualize by CCA in the main text (Figure 6); the full set of KOs was used for all other analyses, as well as the CCA of all KOs in Supplementary Information (Figure S6). Starting with all KOs (5775), KOs were sequentially filtered if they were not found in all samples (3311 remained), if they had a relative count abundance across all samples less than one per thousand counts or 0.001 (2455 remained), or if they had variance less than 5e-8 (252 remained); after CCA, only KOs with non-hypothetical functions were plotted (224 remained). KOs were colored by pathway; note that most KOs belong to multiple KEGG pathways, each with two tiers below ‘Metabolism’. In order to keep the number of pathways displayed to a digestible number, pathways were colored as follows: KOs among the top-ten most represented second-tier pathways were assigned if those pathways had at least five representatives (this corresponded to seven second-tier pathways in Figure 3); the remaining KOs among the top-ten most represented first-tier pathways were assigned if those pathways had at least five representatives (this corresponded to seven first-tier pathways in Figure 3); the remaining KOs were assign to pathway ‘Other’.

### Supplementary Figures

**Figure S1.**
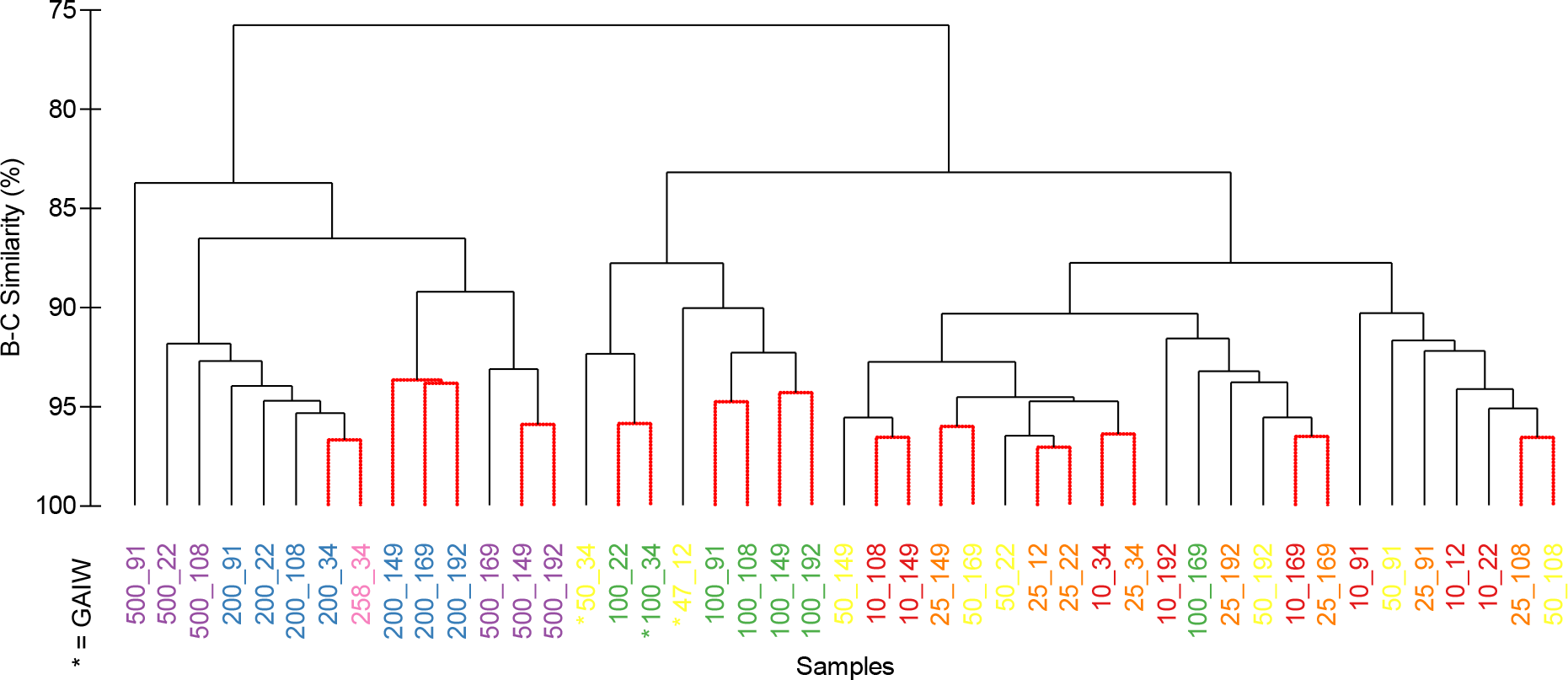
Similarity profile analysis (SIMPROF) of KO relative abundance data using Bray—Curtis similarity. Samples are colored by depth layer, and Gulf of Aden Intermediate Water (GAIW) samples are marked with an asterisk (*).

**Figure S2.**
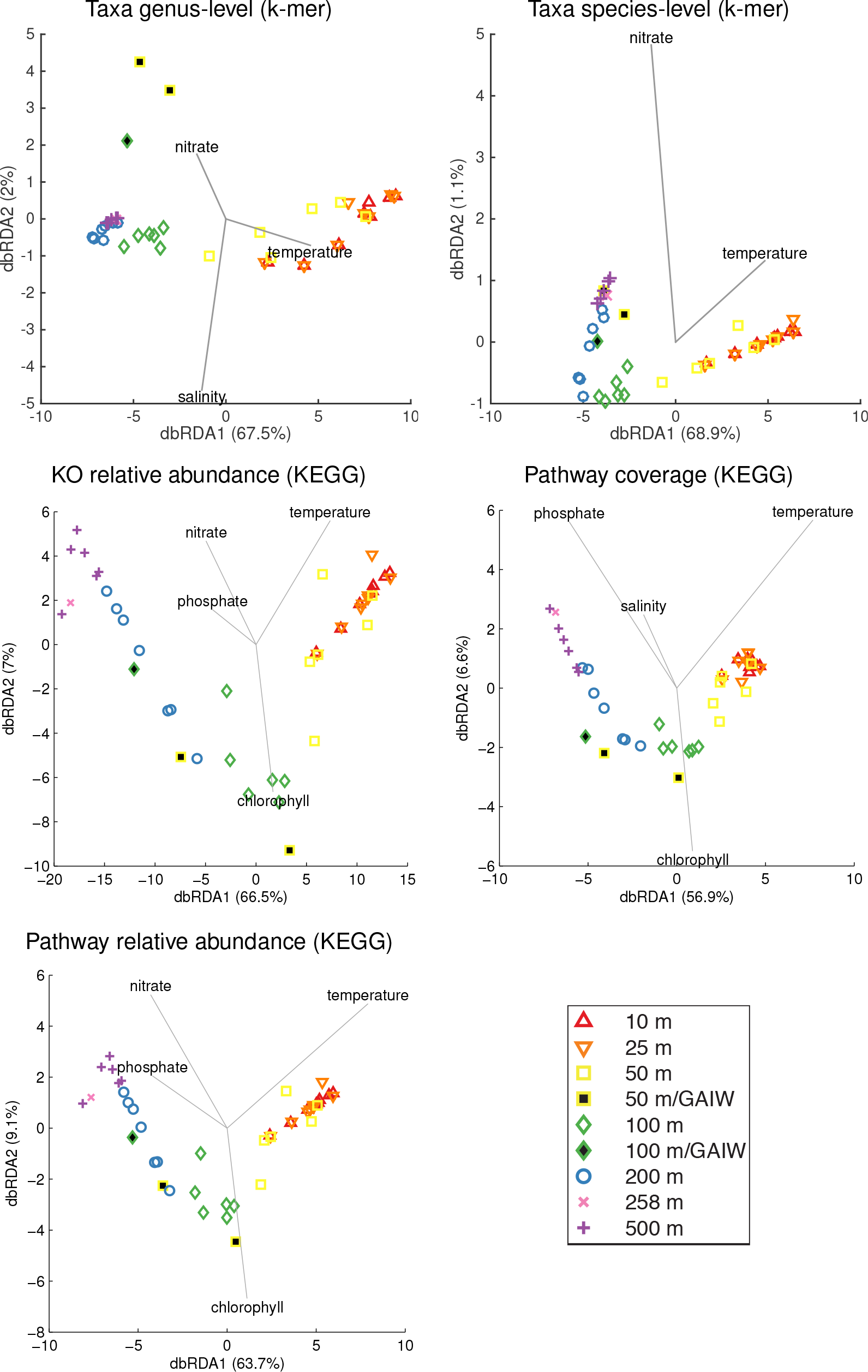
Distance-based redundancy analysis (dbRDA) plots for each response variable. The dbRDA ordination maximizes linear relations of response variables with the predictors. Environmental parameters in the optimal model to the AICc model are plotted. Percent total variation explained by axes 1 and 2 is given.

**Figure S3.**
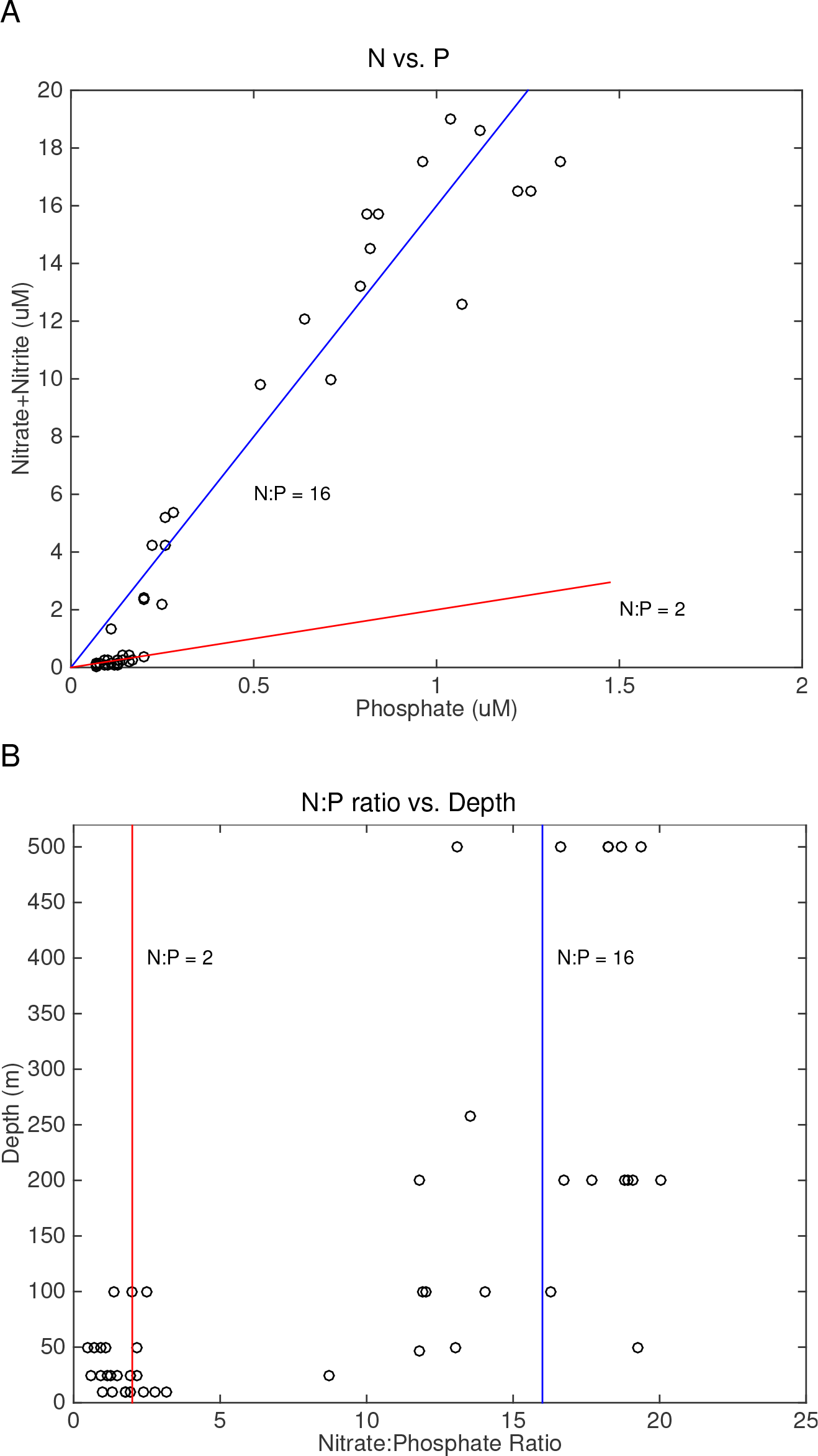
N:P ratio, calculated as nitrate+nitrite to phosphate ratio, across samples plotted as (A) nitrate+nitrite vs. phosphate and (B) depth vs. N:P ratio. Typical Redfield ratio of N:P = 16 is shown as well as the observed N:P = 2 in surface samples.

**Figure S4.**
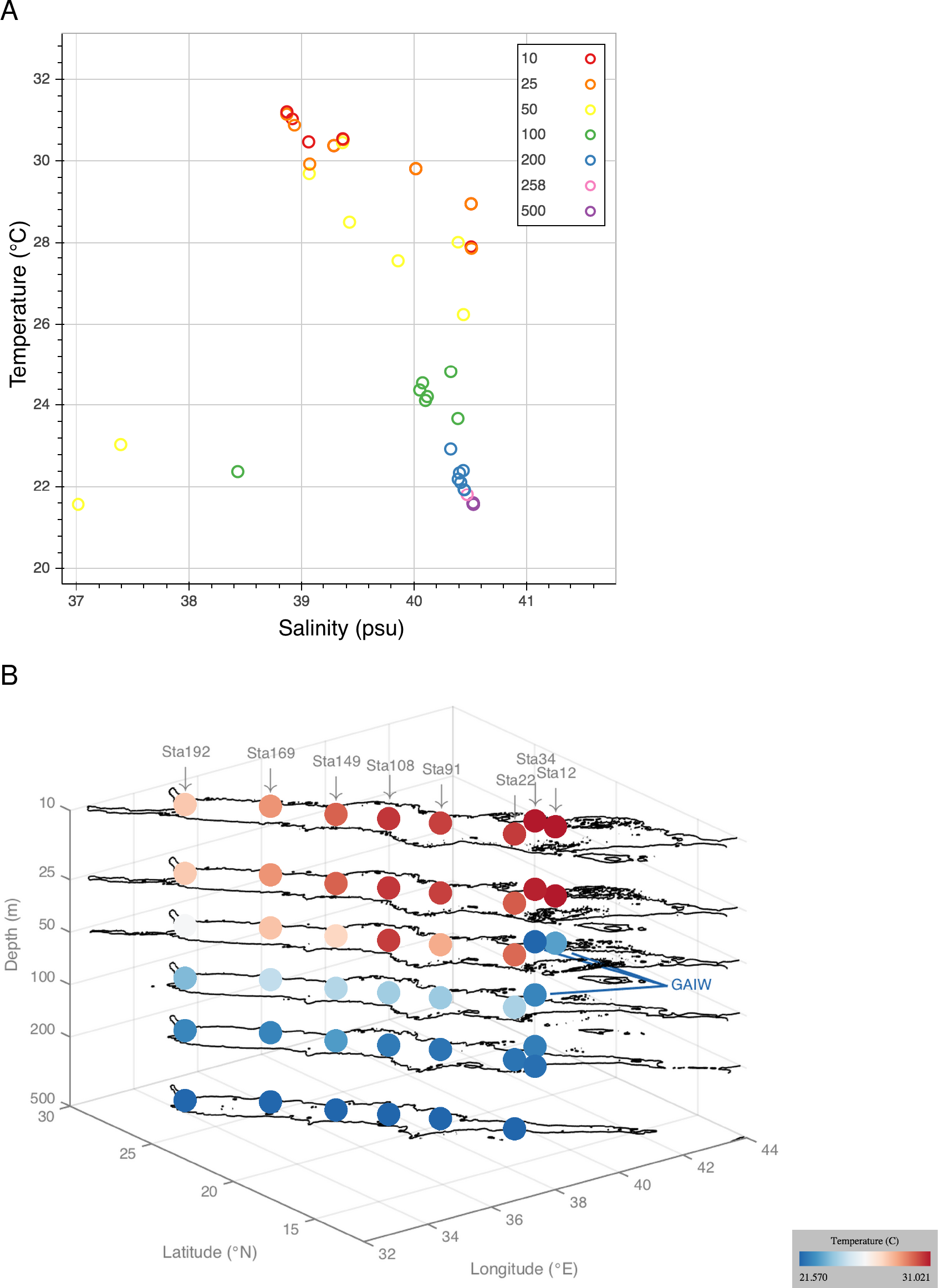
Temperature—salinity (T—S) relationship shown using publicly available interactive visualization tools. (A) T—S profile generated with Bokeh Python package and (B) 3D map of Red Sea colored by temperature generated with ili Toolbox. Points shown are the 45 samples used in this study. The three GAIW foreign water mass samples are clearly visible as distinct from native Red Sea water mass samples by T—S profile and temperature anomaly in the water column.

**Figure S5.**
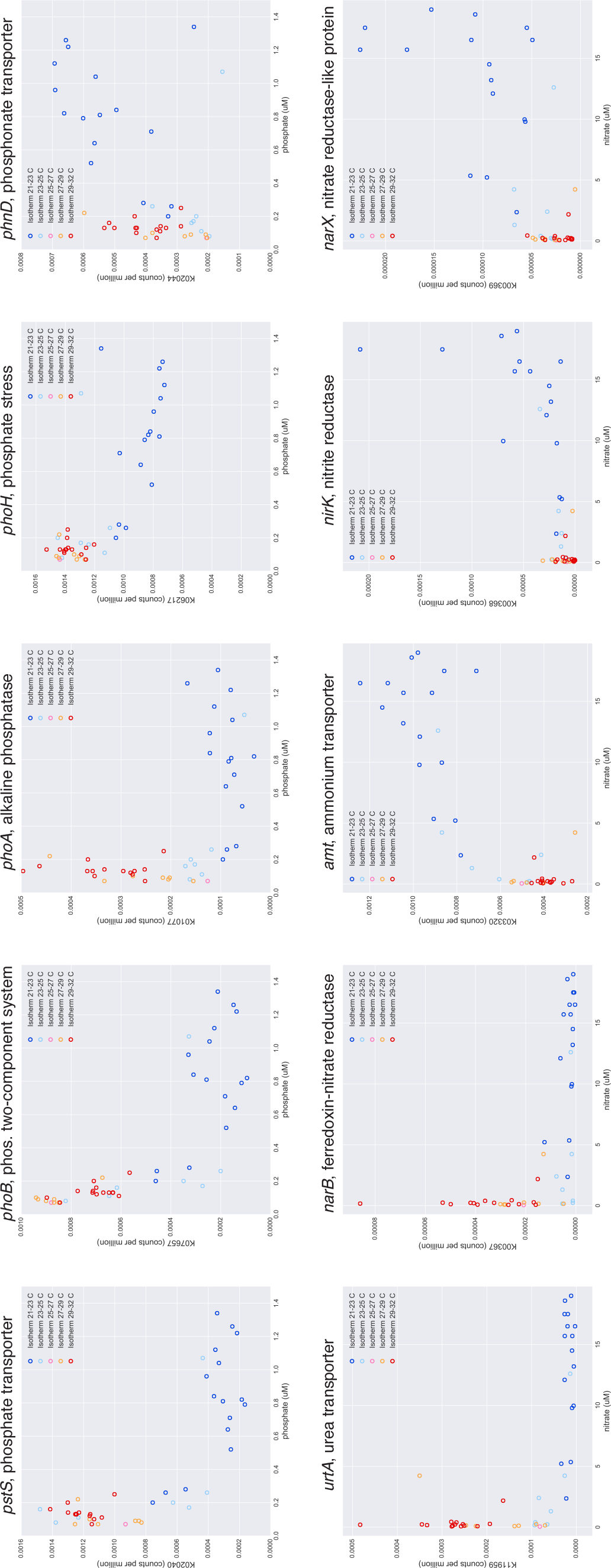
Covariation of nutrient acquisition KOs with phosphate and nitrate, separated by isotherms in 2-degree increments. KO relative abundance is given in units of counts per million of total KO counts in each sample (i.e., all KOs sum to 1 million in each sample).

**Figure S6.**
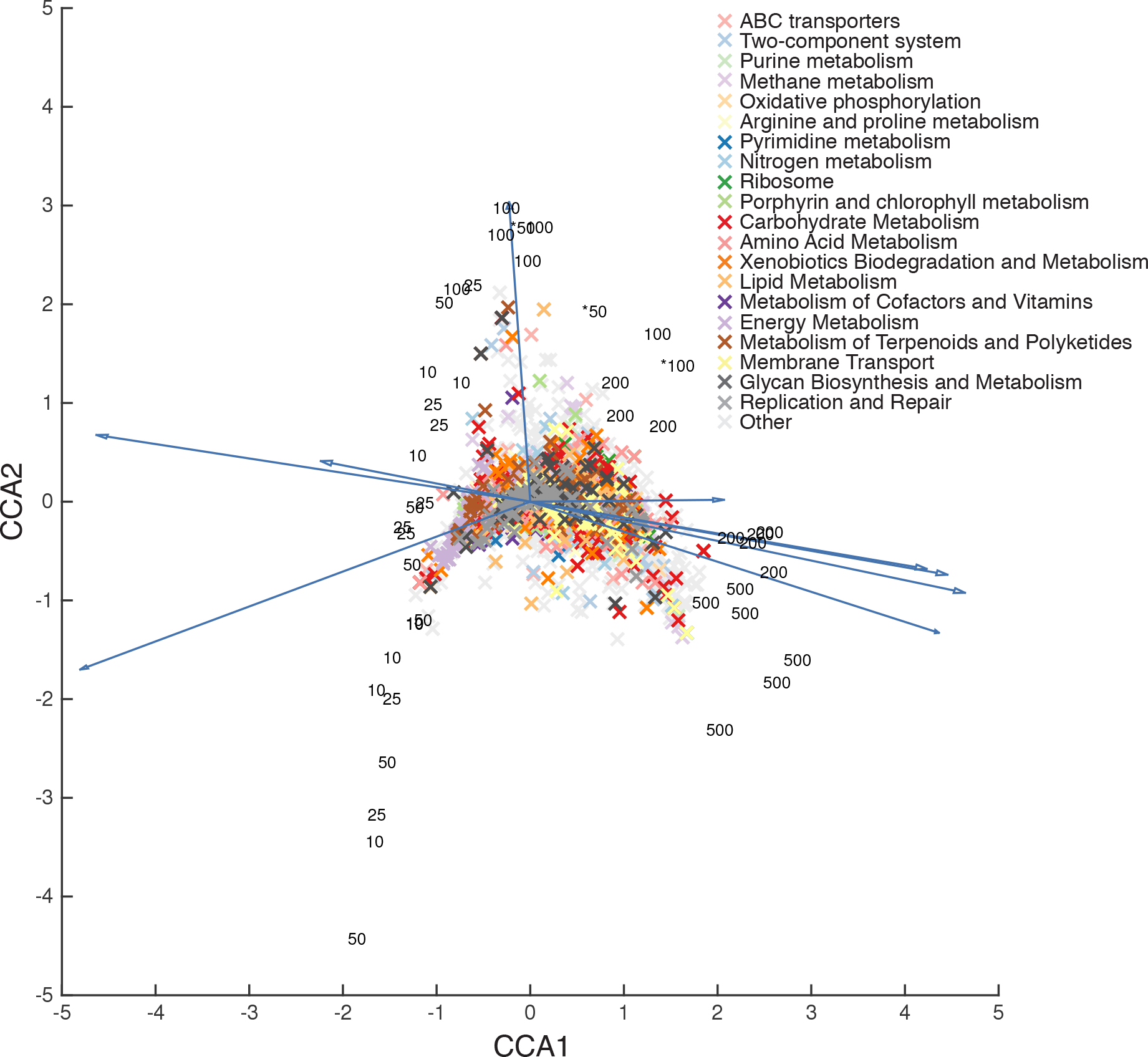
Canonical correspondence analysis of KO relative abundance with environmental parameters, with all KOs displated. Samples are shown as black numerals indicating depth in meters (GAIW samples marked with asterisk), environmental parameters as dark blue arrows, and KOs colored by pathway.

### Supplementary Tables

**Table S1.** Station properties. For each station, the following oceanographic features were calculated from CTD measurements: mixed layer depth (temperature decrease of 0.5 °C from surface), chlorophyll maximum, and oxygen minimum. Values of the chlorophyll maximum and oxygen minimum are given.

| Station | Latitude<br>(°N) | Longitude<br>(°E) | Mixed layer<br>(m) | Chlorophyll max.<br>(m) (mg/m<sup>3</sup>) | Oxygen min.<br>(m) (mL/L) |
| --- | --- | --- | --- | --- | --- |
| 12 | 17.662 | 40.905 | 31 | 44 (2.351) | n/a |
| 22 | 17.996 | 39.799 | 57 | 63 (1.159) | 311 (0.554) |
| 34 | 18.580 | 40.743 | 40 | 48 (1.563) | n/a |
| 91 | 20.525 | 38.781 | 35 | 83 (0.502) | 278 (0.487) |
| 108 | 22.046 | 37.929 | 53 | 97 (0.443) | 381 (0.600) |
| 149 | 23.604 | 37.054 | 52 | 99 (0.456) | 465 (0.756) |
| 169 | 25.772 | 36.116 | 47 | 97 (0.390) | 546 (1.311) |
| 192 | 27.897 | 34.507 | 40 | 58 (0.635) | 498 (1.335) |

**Table S2.**
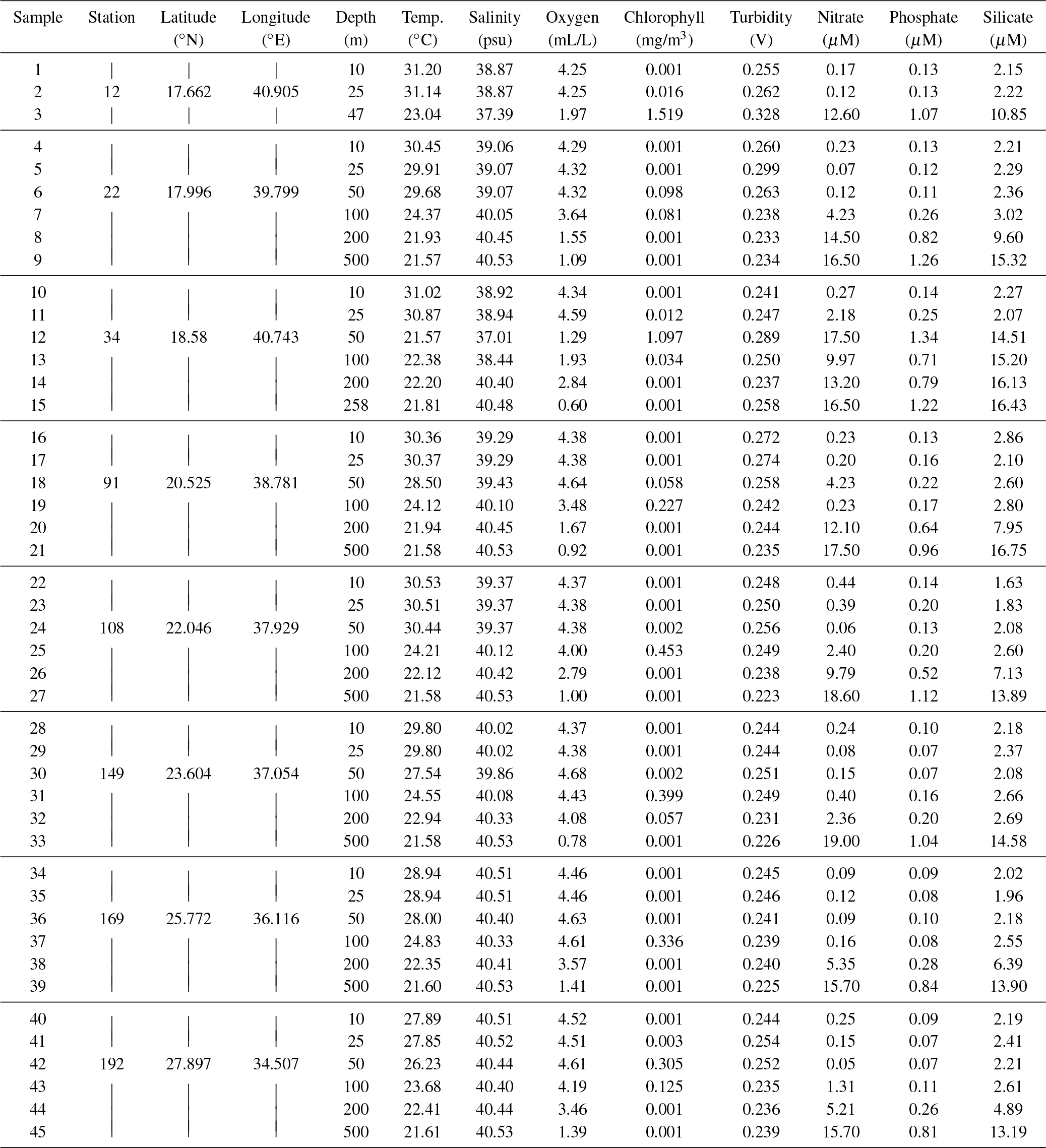
Sample water properties.

**Table S3.** Illumina metagenome properties. Number of reads and total size in bp of forward (Fwd) and reverse (Rev) sequenced reads are after PRINSEQ preprocessing.

| Sample | Station | Insert size |  |  |  | Paired reads<br>(M) | Total basepairs<br>(Gbp) |
| --- | --- | --- | --- | --- | --- | --- | --- |
|  |  | (median, bp) | (MAD, bp) | (mean, bp) | (SD, bp) |  |  |
| 1 | 12 | 299 | 118.6 | 311.6 | 130.8 | 9.6 | 1.8 |
| 2 |  | 305 | 118.6 | 315.5 | 129.7 | 13.5 | 2.5 |
| 3 |  | 284 | 120.1 | 294.2 | 128.1 | 11.5 | 2.1 |
| 4 | 22 | 204 | 106.7 | 215.0 | 102.7 | 9.9 | 1.8 |
| 5 |  | 270 | 115.6 | 278.7 | 121.6 | 7.7 | 1.4 |
| 6 |  | 274 | 120.1 | 282.9 | 124.8 | 7.7 | 1.4 |
| 7 |  | 279 | 117.1 | 288.7 | 124.6 | 8.8 | 1.6 |
| 8 |  | 273 | 117.1 | 282.0 | 122.8 | 8.0 | 1.5 |
| 9 |  | 356 | 129.0 | 367.8 | 140.6 | 8.1 | 1.5 |
| 10 | 34 | 316 | 121.6 | 324.9 | 131.7 | 13.5 | 2.5 |
| 11 |  | 302 | 124.5 | 311.4 | 132.8 | 11.5 | 2.1 |
| 12 |  | 286 | 124.5 | 292.1 | 126.8 | 9.9 | 1.8 |
| 13 |  | 230 | 111.2 | 237.8 | 108.0 | 7.7 | 1.4 |
| 14 |  | 258 | 117.1 | 264.3 | 117.0 | 7.7 | 1.4 |
| 15 |  | 283 | 124.5 | 288.3 | 126.7 | 8.8 | 1.6 |
| 16 | 91 | 359 | 133.4 | 373.3 | 145.7 | 15.7 | 2.9 |
| 17 |  | 366 | 134.9 | 380.1 | 147.9 | 8.1 | 1.5 |
| 18 |  | 244 | 134.9 | 252.5 | 124.8 | 8.0 | 1.5 |
| 19 |  | 186 | 106.7 | 198.8 | 98.4 | 11.5 | 2.1 |
| 20 |  | 183 | 102.3 | 194.8 | 93.8 | 9.9 | 1.8 |
| 21 |  | 233 | 117.1 | 239.4 | 111.2 | 11.3 | 2.1 |
| 22 | 108 | 353 | 129.0 | 362.4 | 139.8 | 11.7 | 2.2 |
| 23 |  | 273 | 123.1 | 279.2 | 123.6 | 10.5 | 2.0 |
| 24 |  | 203 | 102.3 | 211.6 | 97.6 | 11.6 | 2.2 |
| 25 |  | 284 | 126.0 | 289.8 | 126.7 | 9.6 | 1.8 |
| 26 |  | 274 | 127.5 | 279.4 | 125.9 | 11.1 | 2.1 |
| 27 |  | 277 | 124.5 | 283.6 | 125.8 | 10.5 | 2.0 |
| 28 | 149 | 315 | 129.0 | 322.8 | 135.8 | 11.5 | 2.1 |
| 29 |  | 357 | 136.4 | 369.0 | 146.8 | 9.6 | 1.8 |
| 30 |  | 316 | 129.0 | 325.6 | 136.0 | 11.7 | 2.2 |
| 31 |  | 339 | 129.0 | 352.6 | 139.7 | 8.0 | 1.5 |
| 32 |  | 219 | 103.8 | 228.6 | 104.2 | 11.6 | 2.2 |
| 33 |  | 245 | 115.6 | 252.6 | 114.2 | 9.6 | 1.8 |
| 34 | 169 | 328 | 126.0 | 337.4 | 136.3 | 13.5 | 2.5 |
| 35 |  | 284 | 121.6 | 295.1 | 129.6 | 10.5 | 2.0 |
| 36 |  | 305 | 123.1 | 313.1 | 129.2 | 13.9 | 2.6 |
| 37 |  | 227 | 109.7 | 237.8 | 109.5 | 11.3 | 2.1 |
| 38 |  | 245 | 120.1 | 252.4 | 115.6 | 13.5 | 2.5 |
| 39 |  | 188 | 100.8 | 199.4 | 95.3 | 8.8 | 1.6 |
| 40 | 192 | 324 | 124.5 | 334.2 | 133.5 | 15.7 | 2.9 |
| 41 |  | 348 | 126.0 | 357.4 | 136.9 | 8.1 | 1.5 |
| 42 |  | 337 | 127.5 | 344.9 | 136.7 | 13.5 | 2.5 |
| 43 |  | 364 | 132.0 | 372.1 | 141.7 | 10.9 | 2.0 |
| 44 |  | 267 | 121.6 | 273.5 | 120.6 | 11.5 | 2.1 |
| 45 |  | 259 | 115.6 | 266.8 | 117.3 | 11.3 | 2.1 |
|  | Max: | 366 |  | 380.1 |  | Total: 477.7 | 88.8 |
|  | Min: | 183 |  | 194.8 |  |  |  |
|  | Mean: | 282.7 |  | 291.9 |  |  |  |

**Table S4.** Parameters used in PRINSEQ preprocessing, listed in the order that processing steps were applied.

| Parameter | Value | Description |
| --- | --- | --- |
| trim_qual_left | 20 | Trim sequence from 5'-end with quality threshold of 20 |
| trim_qual_right | 20 | Trim sequence from 3'-end with quality threshold of 20 |
| trim_ns_left | 1 | Trim poly-N tail from 5'-end |
| trim_ns_right | 1 | Trim poly-N tail from 3'-end |
| min_len | 40 | Filter sequences shorter than 40 bp (after trimming) |
| min_qual_mean | 20 | Filter sequences with mean quality threshold of 20 (after trimming) |
| ns_max_p | 5 | Filter sequences with more than 5% Ns (after trimming) |
| lc_method | entropy | Filter low complexity sequences with Shannon entropy... |
| lc_threshold | 50 | ... less than 50 (after trimming) |
| derep | 14 | Filter exact duplicates and reverse complement exact duplicates (after trimming) |

**Table S5.** Results of AICc, the stepwise explanation of variation in response variables by sequentially adding environmental parameters (predictors), balancing performance and parsimony.

File: Table_S5_AICc_Results.xlsx

**Table S6.** Percent of reads mapped by HUMAnN and taxonomy assignment methods.Columns from left to right: HUMAnN translated search to prokaryotic KO sequences in KEGG; CLARK genus-level k-mer; CLARK species-level k-mer; Kraken genus-level k-mer; GraftM genus-level 16S; GraftM ecotype-level *Prochlorococcus rpoC1*; GraftM ecotype-level *Pelagibacter* 16S.

File: Table_S6_Percent_Reads_Mapped.xlsx

**Table S7.** KOs ranked by total abundance across all samples.

File: Table_S7_KOs_Ranked_By_Abundance.xlsx

**Table S8.** KOs clustered by total abundance across all samples using partitioning around medoids (PAM) with 12 clusters.

File: Table_S8_KOs_Partitioned_Around_Medoids.xlsx

**Table S9.**
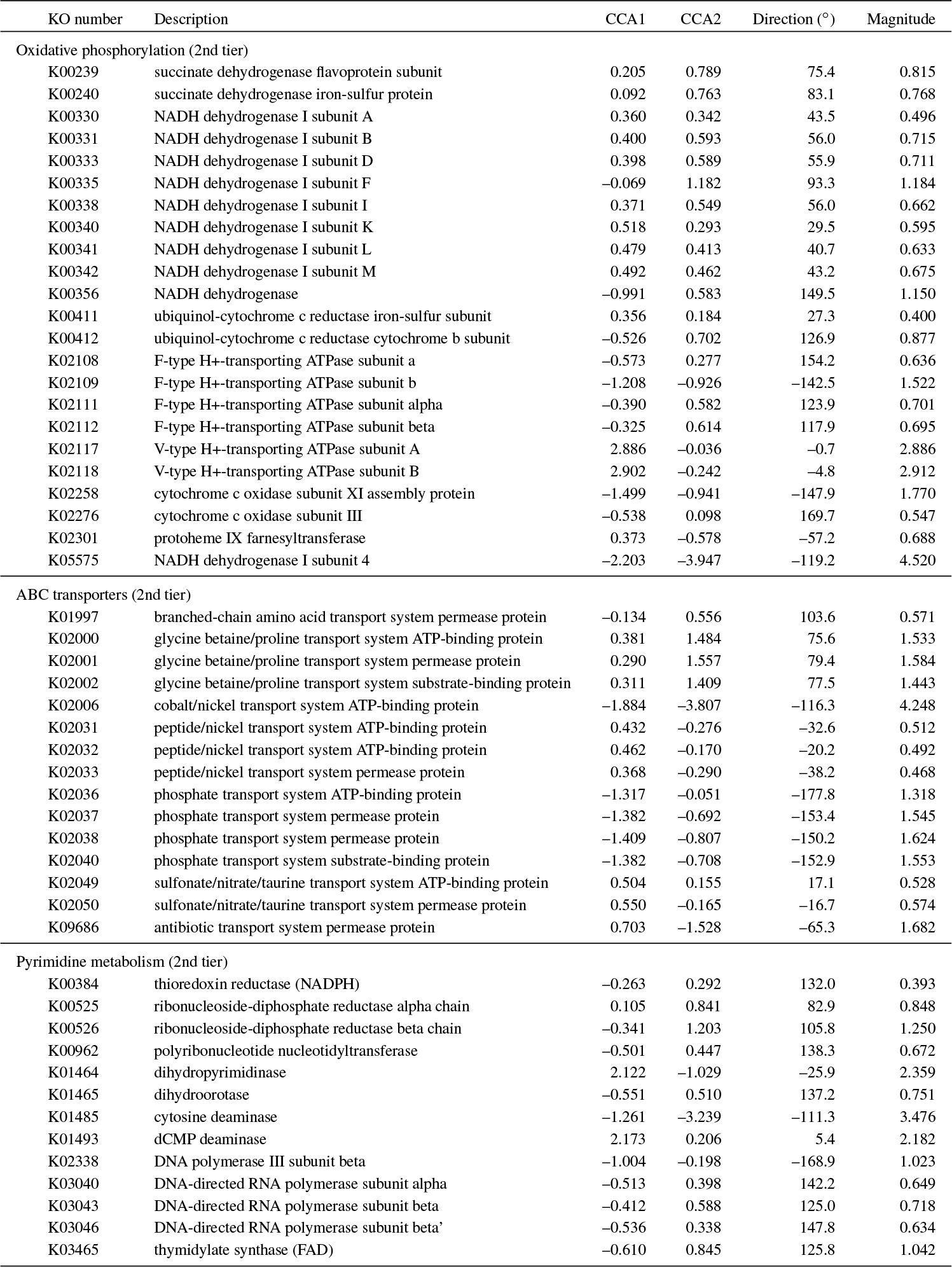
Cartesian and polar coordinates of the 224 KEGG KOs (gene ortholog groups) plotted in the CCA ordination. KOs were first assigned to second-tier KEGG pathways (if total pathway count was at least 5), then first-tier KEGG pathways (if total pathway count was at least 5), and the remaining KOs were grouped as ‘Other’. Direction is in degrees from the polar axis. For reference, environmental parameter directions are as follows: salinity, 0.5°; chlorophyll, 94.3°; turbidity, 169.6°; oxygen, 171.7°; temperature, −160.5°; depth, −16.9°; nitrate, −11.2°; silicate, −9.4°; phosphate, −9.1°.

| KO number | Description | CCA1 | CCA2 | Direction (°) | Magnitude |
| --- | --- | --- | --- | --- | --- |
| Oxidative phosphorylation (2nd tier) |  |  |  |  |  |
| K00239 | succinate dehydrogenase flavoprotein subunit | 0.205 | 0.789 | 75.4 | 0.815 |
| K00240 | succinate dehydrogenase iron-sulfur protein | 0.092 | 0.763 | 83.1 | 0.768 |
| K00330 | NADH dehydrogenase I subunit A | 0.360 | 0.342 | 43.5 | 0.496 |
| K00331 | NADH dehydrogenase I subunit B | 0.400 | 0.593 | 56.0 | 0.715 |
| K00333 | NADH dehydrogenase I subunit D | 0.398 | 0.589 | 55.9 | 0.711 |
| K00335 | NADH dehydrogenase I subunit F | -0.069 | 1.182 | 93.3 | 1.184 |
| K00338 | NADH dehydrogenase I subunit I | 0.371 | 0.549 | 56.0 | 0.662 |
| K00340 | NADH dehydrogenase I subunit K | 0.518 | 0.293 | 29.5 | 0.595 |
| K00341 | NADH dehydrogenase I subunit L | 0.479 | 0.413 | 40.7 | 0.633 |
| K00342 | NADH dehydrogenase I subunit M | 0.492 | 0.462 | 43.2 | 0.675 |
| K00356 | NADH dehydrogenase | -0.991 | 0.583 | 149.5 | 1.150 |
| K00411 | ubiquinol-cytochrome c reductase iron-sulfur subunit | 0.356 | 0.184 | 27.3 | 0.400 |
| K00412 | ubiquinol-cytochrome c reductase cytochrome b subunit | -0.526 | 0.702 | 126.9 | 0.877 |
| K02108 | F-type H<sup>+</sup>-transporting ATPase subunit a | -0.573 | 0.277 | 154.2 | 0.636 |
| K02109 | F-type H<sup>+</sup>-transporting ATPase subunit b | -1.208 | -0.926 | -142.5 | 1.522 |
| K02111 | F-type H<sup>+</sup>-transporting ATPase subunit alpha | -0.390 | 0.582 | 123.9 | 0.701 |
| K02112 | F-type H<sup>+</sup>-transporting ATPase subunit beta | -0.325 | 0.614 | 117.9 | 0.695 |
| K02117 | V-type H<sup>+</sup>-transporting ATPase subunit A | 2.886 | -0.036 | -0.7 | 2.886 |
| K02118 | V-type H<sup>+</sup>-transporting ATPase subunit B | 2.902 | -0.242 | -4.8 | 2.912 |
| K02258 | cytochrome c oxidase subunit XI assembly protein | -1.499 | -0.941 | -147.9 | 1.770 |
| K02276 | cytochrome c oxidase subunit III | -0.538 | 0.098 | 169.7 | 0.547 |
| K02301 | protoheme IX farnesyltransferase | 0.373 | -0.578 | -57.2 | 0.688 |
| K05575 | NADH dehydrogenase I subunit 4 | -2.203 | -3.947 | -119.2 | 4.520 |
| ABC transporters (2nd tier) |  |  |  |  |  |
| K01997 | branched-chain amino acid transport system permease protein | -0.134 | 0.556 | 103.6 | 0.571 |
| K02000 | glycine betaine/proline transport system ATP-binding protein | 0.381 | 1.484 | 75.6 | 1.533 |
| K02001 | glycine betaine/proline transport system permease protein | 0.290 | 1.557 | 79.4 | 1.584 |
| K02002 | glycine betaine/proline transport system substrate-binding protein | 0.311 | 1.409 | 77.5 | 1.443 |
| K02006 | cobalt/nickel transport system ATP-binding protein | -1.884 | -3.807 | -116.3 | 4.248 |
| K02031 | peptide/nickel transport system ATP-binding protein | 0.432 | -0.276 | -32.6 | 0.512 |
| K02032 | peptide/nickel transport system ATP-binding protein | 0.462 | -0.170 | -20.2 | 0.492 |
| K02033 | peptide/nickel transport system permease protein | 0.368 | -0.290 | -38.2 | 0.468 |
| K02036 | phosphate transport system ATP-binding protein | -1.317 | -0.051 | -177.8 | 1.318 |
| K02037 | phosphate transport system permease protein | -1.382 | -0.692 | -153.4 | 1.545 |
| K02038 | phosphate transport system permease protein | -1.409 | -0.807 | -150.2 | 1.624 |
| K02040 | phosphate transport system substrate-binding protein | -1.382 | -0.708 | -152.9 | 1.553 |
| K02049 | sulfonate/nitrate/taurine transport system ATP-binding protein | 0.504 | 0.155 | 17.1 | 0.528 |
| K02050 | sulfonate/nitrate/taurine transport system permease protein | 0.550 | -0.165 | -16.7 | 0.574 |
| K09686 | antibiotic transport system permease protein | 0.703 | -1.528 | -65.3 | 1.682 |
| Pyrimidine metabolism (2nd tier) |  |  |  |  |  |
| K00384 | thioredoxin reductase (NADPH) | -0.263 | 0.292 | 132.0 | 0.393 |
| K00525 | ribonucleoside-diphosphate reductase alpha chain | 0.105 | 0.841 | 82.9 | 0.848 |
| K00526 | ribonucleoside-diphosphate reductase beta chain | -0.341 | 1.203 | 105.8 | 1.250 |
| K00962 | polyribonucleotide nucleotidyltransferase | -0.501 | 0.447 | 138.3 | 0.672 |
| K01464 | dihydropyrimidinase | 2.122 | -1.029 | -25.9 | 2.359 |
| K01465 | dihydroorotase | -0.551 | 0.510 | 137.2 | 0.751 |
| K01485 | cytosine deaminase | -1.261 | -3.239 | -111.3 | 3.476 |
| K01493 | dCMP deaminase | 2.173 | 0.206 | 5.4 | 2.182 |
| K02338 | DNA polymerase III subunit beta | -1.004 | -0.198 | -168.9 | 1.023 |
| K03040 | DNA-directed RNA polymerase subunit alpha | -0.513 | 0.398 | 142.2 | 0.649 |
| K03043 | DNA-directed RNA polymerase subunit beta | -0.412 | 0.588 | 125.0 | 0.718 |
| K03046 | DNA-directed RNA polymerase subunit beta' | -0.536 | 0.338 | 147.8 | 0.634 |
| K03465 | thymidylate synthase (FAD) | -0.610 | 0.845 | 125.8 | 1.042 |

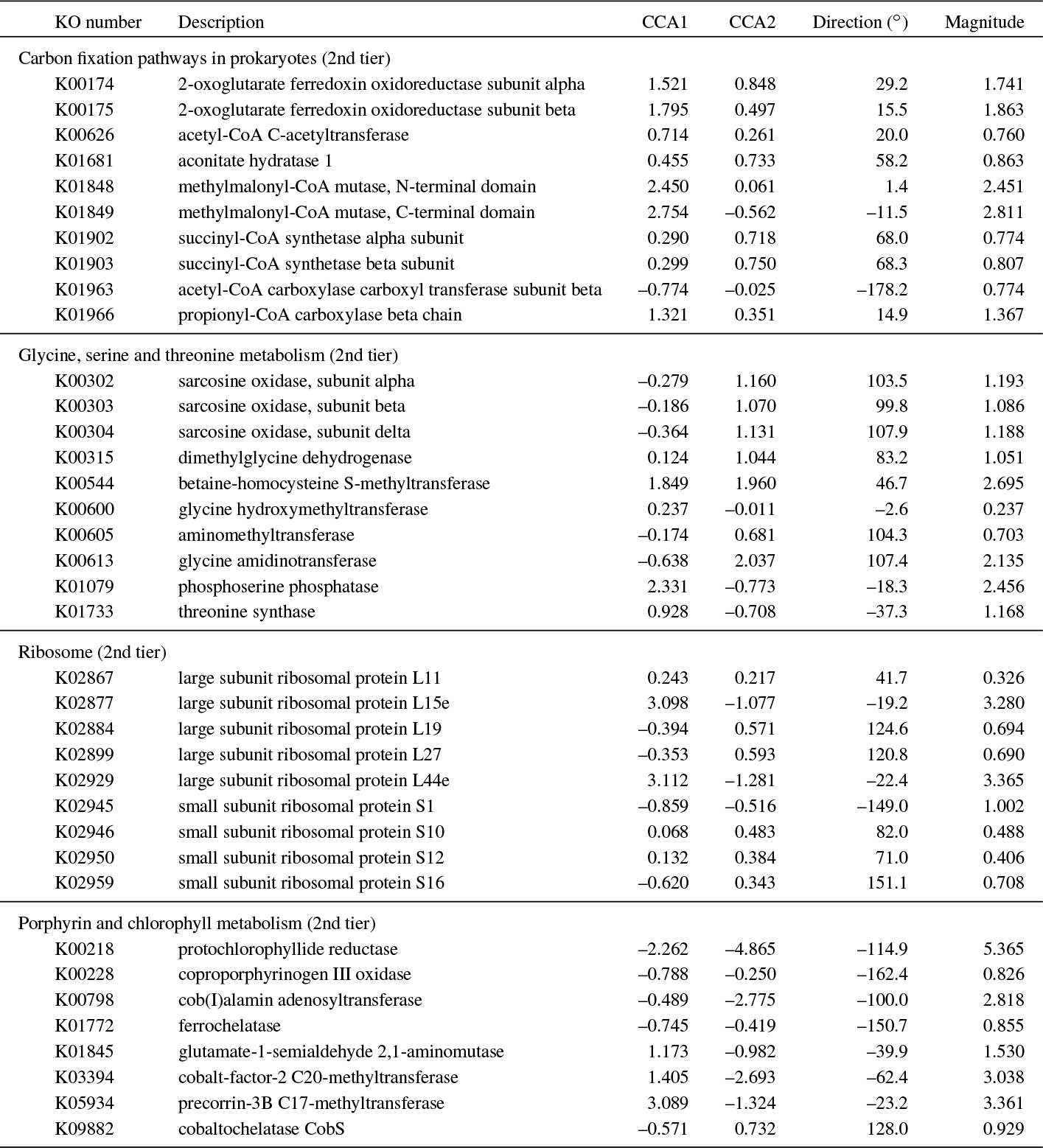

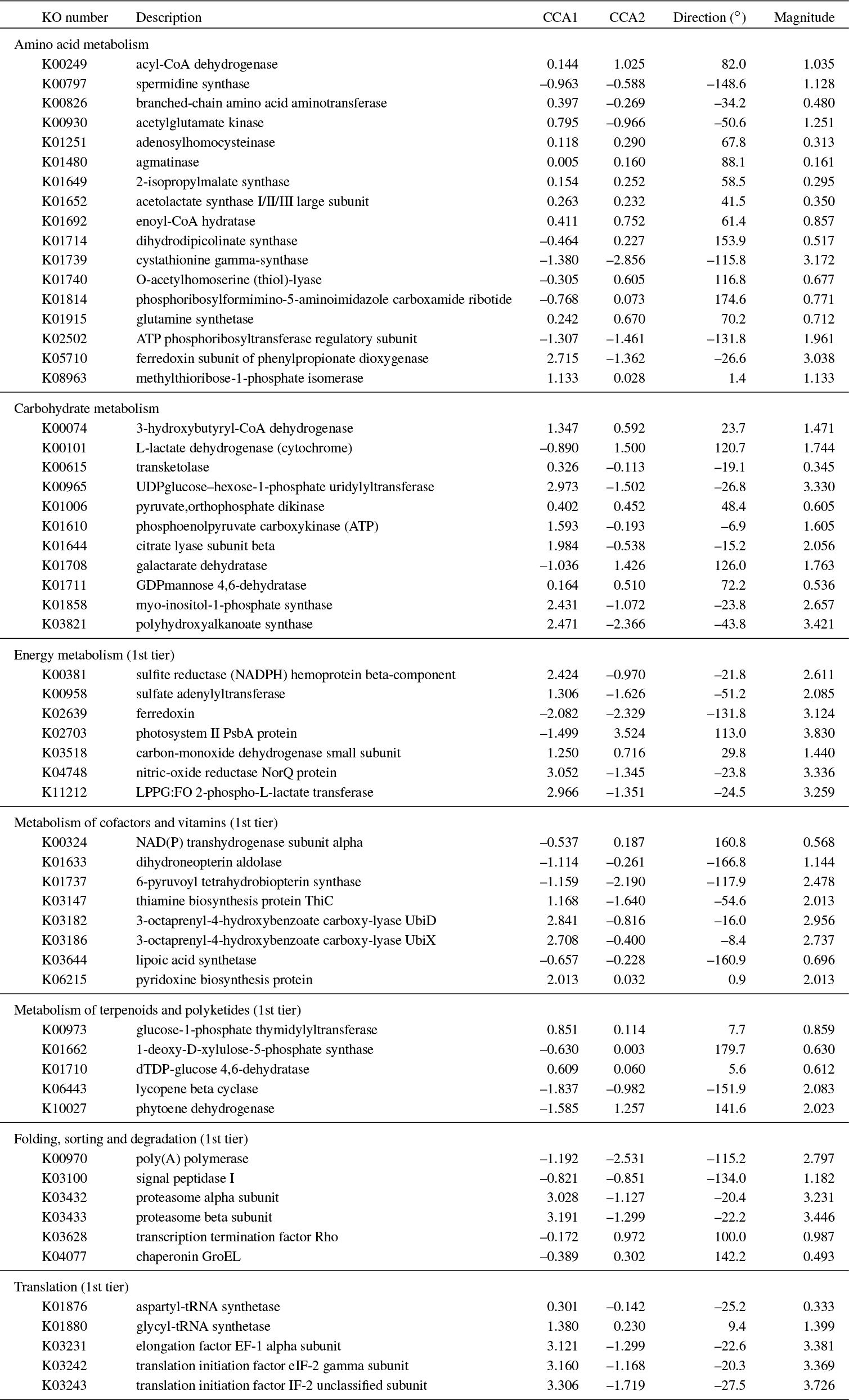

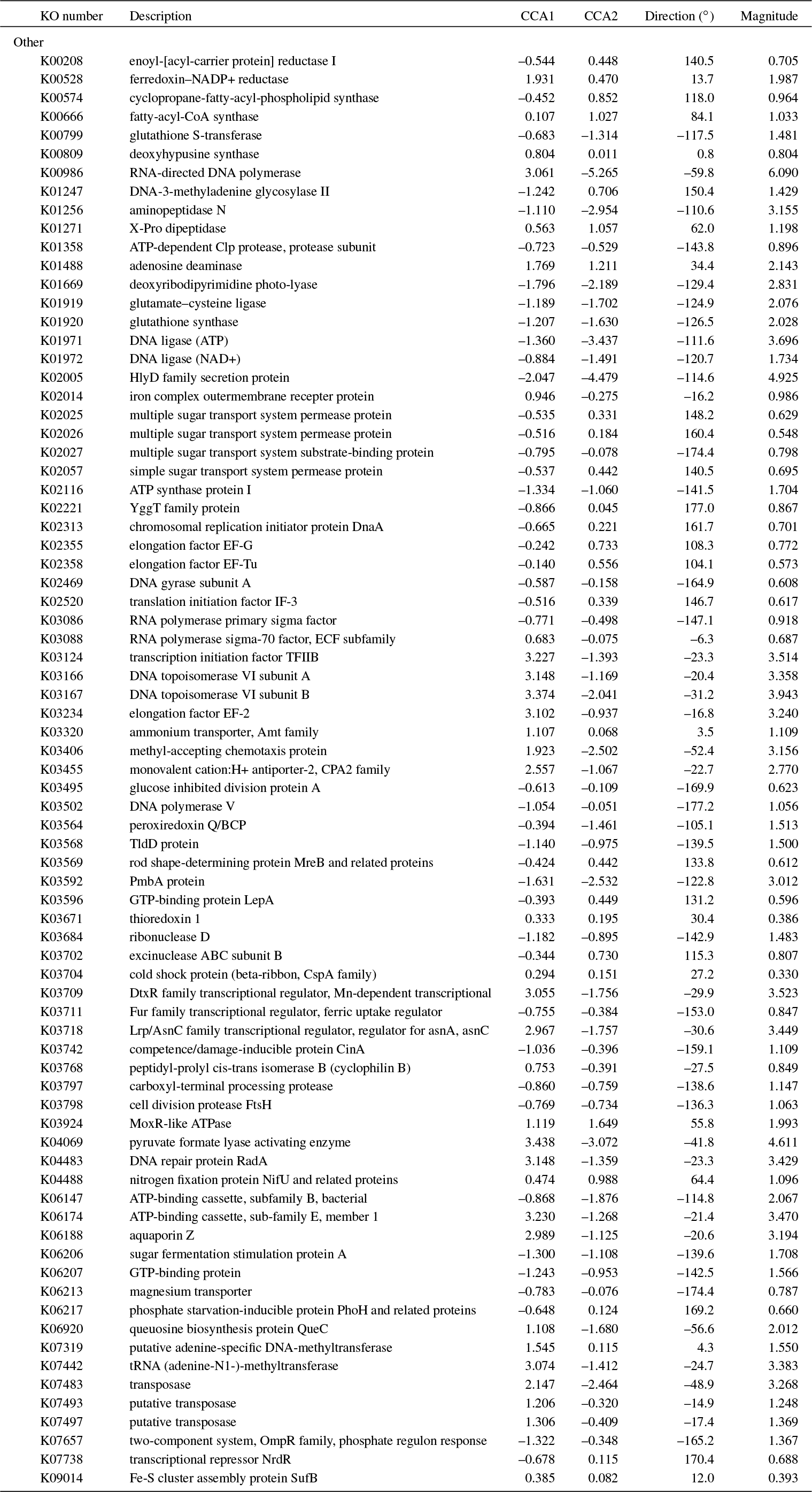

### Online Content

**Interactive 3D maps.** Point the Chrome browser to http://ili-toolbox.github.io/ or install the ili Toolbox Chrome app. Drag the PNG file then one of the CSV data files to the web page. Use the on-screen menu to change the heatmap appearance; we recommend setting spot border to 1, hotspot quantile to 1, and color map to Blue-Red. Select data categories to view or search for terms in the selection menu. Drag another CSV file to change datasets.

Map file (PNG) and data files (CSV) may be downloaded from GitHub:

RedSea_Map.png
RedSea_TaxaRelAbund_Genus.csv
RedSea_TaxaRelAbund_Species.csv
RedSea_TaxaCounts_Pelagibacter.csv
RedSea_TaxaCounts_Prochlorococcus.csv
RedSea_KORelativeAbundance_AllTaxa.csv
RedSea_KORelativeAbundance_Nitrosopumilus.csv
RedSea_KORelativeAbundance_Pelagibacter.csv
RedSea_KORelativeAbundance_Prochlorococcus.csv
RedSea_PathwayCoverage_AllTaxa.csv
RedSea_PathwayRelativeAbundance_AllTaxa.csv
RedSea_PredictedRelativeMetabolicTurnover_AllTaxa.csv

**Interactive scatter plots.** Open the HTML files in any web browser and choose any set of two environmental parameters or response variables to plot. Type the first few letters to go directly to a selection. Plots can be resized and then saved to PNG.

Plot files (HTML) may be downloaded from GitHub:

RedSea_TaxaRelAbund_Genus.html
RedSea_TaxaRelAbund_Species.html
RedSea_TaxaCounts_Pelagibacter.html
RedSea_TaxaCounts_Prochlorococcus.html
RedSea_KORelativeAbundance_AllTaxa.html
RedSea_KORelativeAbundance_Nitrosopumilus.html
RedSea_KORelativeAbundance_Pelagibacter.html
RedSea_KORelativeAbundance_Prochlorococcus.html
RedSea_PathwayCoverage_AllTaxa.html
RedSea_PathwayRelativeAbundance_AllTaxa.html
RedSea_PredictedRelativeMetabolicTurnover_AllTaxa.html

